# HNF4α controls growth, identity and response to KRAS inhibition of invasive mucinous adenocarcinoma of the lung

**DOI:** 10.1101/2025.06.06.658155

**Authors:** Headtlove Essel Dadzie, Yangsook Song Green, Soledad Camolotto, Matthew Gumbleton, Yutaka Maeda, Benjamin T. Spike, Eric L. Snyder

## Abstract

Cellular plasticity is a hallmark of cancer, enabling tumor cells to alter identity and evade therapeutic pressure. In invasive mucinous adenocarcinoma of the lung (IMA), NKX2-1 loss triggers a pulmonary to gastric switch marked by aberrant activation of HNF4α, a master regulator of gastrointestinal/hepatic differentiation. We find that HNF4α promotes IMA growth and activates a gastric pit cell-like program. *Hnf4a* deletion induces IMA dedifferentiation, enabling FoxA1/2 to access de novo sites and activate alternative identities. HNF4α also induces a mucinous program associated with tolerance to KRAS blockade, and HNF4α loss enhances response to KRAS^G12D^ inhibition. Mechanistically, HNF4α blocks cell cycle exit in drug-tolerant persister cells and promotes activity of the antioxidant transcription factor NRF2. NRF2 activation partially rescues effects of *Hnf4a* deletion on KRAS^G12D^ inhibition, whereas NRF2 inhibition enhances sensitivity to KRAS^G12D^ blockade. Thus, HNF4α is a key regulator of identity and primary response to KRAS^G12D^ inhibition in IMA.

**SIGNIFICANCE:** IMA is a genetically and epigenetically distinct LUAD subtype for which targeted therapies are lacking due to the high proportion of KRAS mutations. This study points to blockade of the HNF4α -> NRF2 axis as a potential strategy to enhance primary response to KRAS inhibition in IMA.

## INTRODUCTION

Cancer remains a leading cause of mortality worldwide, driven by its inherent molecular and cellular heterogeneity. Despite significant advances in molecular profiling and targeted therapies, treatment outcomes remain suboptimal across multiple cancer types(Siegel et al., 2025; Tan & Tan, 2022). A key challenge in oncology is the remarkable plasticity of cancer cells, which enables dynamic shifts in cellular identity in response to genetic, epigenetic, and transcriptional cues(Hanahan, 2022; Junttila & de Sauvage, 2013; Tavernari et al., 2021). This plasticity limits the durability of therapeutic response, promoting tumor progression, metastasis and recurrence(Flavahan et al., 2017; Laisné et al., 2025). Understanding the mechanisms that govern cancer cell identity and plasticity is therefore critical for developing more effective treatment strategies.

Lung adenocarcinoma (LUAD), the most prevalent subtype of lung cancer(Siegel et al., 2025), exemplifies these challenges due to its molecular heterogeneity and lineage plasticity(Han et al., 2024; Karasaki et al., 2023). Although therapies targeting driver oncogenes such as EGFR(Ayati et al., 2020), KRAS (particularly G12C)(Isermann et al., 2024; Liu et al., 2022), and ALK(Sullivan & Planchard, 2016) have transformed clinical care, resistance remains common(Awad et al., 2021; Gainor et al., 2016). LUAD cells can transition between epithelial and alternative identities, including neuroendocrine, squamous, or gastric-like states, particularly under therapeutic pressure(Araujo et al., 2024; Camolotto et al., 2018; Davies et al., 2023; Hanahan, 2022). These transitions, driven by epigenetic and transcriptional reprogramming, increase intratumoral heterogeneity and pose barriers to effective treatment(de Visser & Joyce, 2023; Flavahan et al., 2017; Yang et al., 2023).

Invasive mucinous adenocarcinoma (IMA) is a distinct LUAD subtype (∼5-10% of cases)(Kunii et al., 2011) that provides a compelling model to study transcription factor-driven lineage plasticity(Snyder et al., 2013). IMA arises through loss of NKX2-1/TTF1, a master regulator of pulmonary epithelial identity, and acquisition of gastric identity via FoxA1/2-mediated transcriptional reprogramming(Camolotto et al., 2018; Iwafuchi-Doi et al., 2016; Orstad et al., 2022; Snyder et al., 2013). The adult gastric epithelium consists of multiple specialized cell types (e.g., pit, neck, chief, parietal, tuft, and enteroendocrine) that are predominantly derived from isthmus progenitors at steady state(Karam & Leblond, 1993; Stengel & Taché, 2009). IMA tumor cells partially recapitulate gastric differentiation, including a major pit-like population with abundant mucin production as well as a minor tuft-like population(Sugano et al., 2013; Travis et al., 2011; Zewdu et al., 2021). Tuft cells are rare chemosensory sentinels with emerging roles in cancer(DelGiorno et al., 2020). Tuft-like transcriptional states have also been described in epithelial malignancies including colorectal, pancreatic, lung, and thymic tumors(Nakanishi et al., 2013; Yamada et al., 2021; Yamashita et al., 2017). We have previously shown that levels of MEK/ERK and WNT signaling dictate the specific gastric cell types that IMA cells resemble(Zewdu et al., 2021), which is consistent with the effects of these signaling pathways on differentiation in the normal stomach(Takada et al., 2023).

The nuclear receptor HNF4α(Tirona et al., 2003) has emerged as a candidate regulator of gastric lineage fidelity in IMA(Snyder et al., 2013). HNF4α is abundantly expressed in liver and GI epithelia where it regulates metabolic and differentiation programs(Chandra et al., 2013; Jiang et al., 2003; Lau et al., 2018; Moore et al., 2016). Notably, HNF4α is absent in normal lung tissue, but robustly induced by FoxA1/2 in IMA(Camolotto et al., 2018). Depending on cellular context, HNF4α has been reported to act as a tumor suppressor or oncogene(Camolotto et al., 2021; Chellappa et al., 2016; Lv et al., 2021; Xu et al., 2001; Yin et al., 2008), contributing to tumor growth, metastasis, and resistance to apoptosis(Walesky et al., 2013; Xu et al., 2020).

Up to 75% of IMA harbor KRAS mutations(Prior et al., 2020), with the G12D variant being the most common(Chang et al., 2021). Targeted therapies against KRAS^G12D^ are currently under active preclinical and clinical evaluation in multiple cancers(Hallin et al., 2022; Knox et al., 2022; Y. Li et al., 2025). However, mechanisms of primary response and secondary resistance to KRAS inhibition remain incompletely understood. The antioxidant transcription factor NRF2 is emerging as a modulator of response to KRAS inhibition in LUAD(Negrao et al., 2025). Under basal conditions, NRF2 is targeted for degradation by KEAP1, but oxidative stress or inactivating *KEAP1* mutation leads to NRF2 stabilization(Ulasov et al., 2022). NRF2 activity has been associated with resistance to KRAS inhibitors even in functionally KEAP1-wildtype LUAD, suggesting non-genetic mechanisms of NRF2 activation(Lignitto et al., 2019; Negrao et al., 2025; Romero et al., 2017). Recent evidence also identifies NRF2 as a direct transcriptional target of HNF4α in renal epithelial cells, raising the possibility that HNF4α might contribute to NRF2 activation and therapeutic resistance in IMA(X. Li et al., 2025).

Using a genetically engineered mouse model (GEMM) that recapitulates key features of human IMA, we previously showed that *Hnf4a* deletion partially impairs the IMA initiation(Snyder et al., 2013). Studies in human cell lines have identified potential mechanisms by which HNF4α might support the IMA growth(Chen et al., 2021; Stuart et al., 2024). HNF4α and other GI markers have recently been shown to be enriched in drug-tolerant persister cells (DTPs) in LUAD GEMMs treated with RAS inhibitors(Araujo et al., 2024). However, critical mechanistic questions remain: 1) Is HNF4α required for the maintenance of established IMA growth in vivo? 2) What is the interplay between HNF4α and other lineage specifiers, such as FoxA1/2, in controlling IMA identity? 3) Does HNF4α modulate therapeutic response to KRAS inhibition, or does its expression merely correlate with persistence? Here, we address these questions using IMA GEMMs, mouse- and patient-derived organoids (PDOs) and integrative multi-omics approaches(Stuart & Satija, 2019) to uncover the role of HNF4α in IMA lineage fidelity and response to targeted therapy.

## RESULTS

### HNF4α is essential for in vivo growth of established IMA

To investigate the role of HNF4α in established IMA tumors in vivo, we developed an advanced GEMM model utilizing sequential activation of Flp and Cre recombinases for the conditional deletion of *Hnf4a* in established tumors (**Sup. Fig. S1A**). Tumors were initiated via intratracheal administration of an adenovirus carrying codon-optimized Flp recombinase (FlpO) under the control of the surfactant protein C gene (*SPC*) promoter(Camolotto et al., 2018). This system activates *Kras^G12D^* and *Cre^ERT2^* transcription in distal lung epithelial cells(Schönhuber et al., 2014; Young et al., 2011).

Tamoxifen treatment of this mouse model induced efficient Cre^ERT2^-mediated recombination in established lesions, leading to the near complete deletion of *Nkx2-1*, in tumors harboring wild type *Hnf4a* alleles (*Kras^FSF-G12D/+^*; *Rosa26^FSF-CreERT2^; Nkx2-1^F/F^; Hnf4a^+/+^* or *Kras^FSF-G12D/+^*; *Rosa26^FSF-CreERT2^; Nkx2-1^F/F^; Hnf4a^F/+^*, hereafter referred to as KN) as well as in those with homozygous deletion of floxed *Hnf4a* alleles (*Kras^FSF-G12D/+^*; *Rosa26^FSF-CreERT2^; Nkx2-1^F/F^; Hnf4a^F/F^*, *designated as* KNH) (**Fig. 1A**). Morphologically, KNH tumor cells exhibited an increased nuclear-to-cytoplasmic ratio and reduced cytoplasmic volume compared to control KN tumor cells (**Fig. 1A**, assessed by board-certified pathologist E.L.S.). At 14 weeks post-tumor initiation (PTI), KNH mice exhibited ∼50% lower tumor burden compared to KN controls (p < 0.001; **Fig. 1B**). Because there were no differences in morphology or tumor burden between *Hnf4a^+/+^* and *Hnf4a^F/+^* KN tumors, we used *Hnf4a^F/+^* KN mice as controls in subsequent experiments. In KNH mice, *Hnf4a* deletion led to increased apoptosis as early as 1-week after the first tamoxifen dose and persisted for 8 weeks (**Fig. 1C**), but did not significantly affect the proliferation rate (**Sup. Fig. S1D-E**).

**Figure 1.**
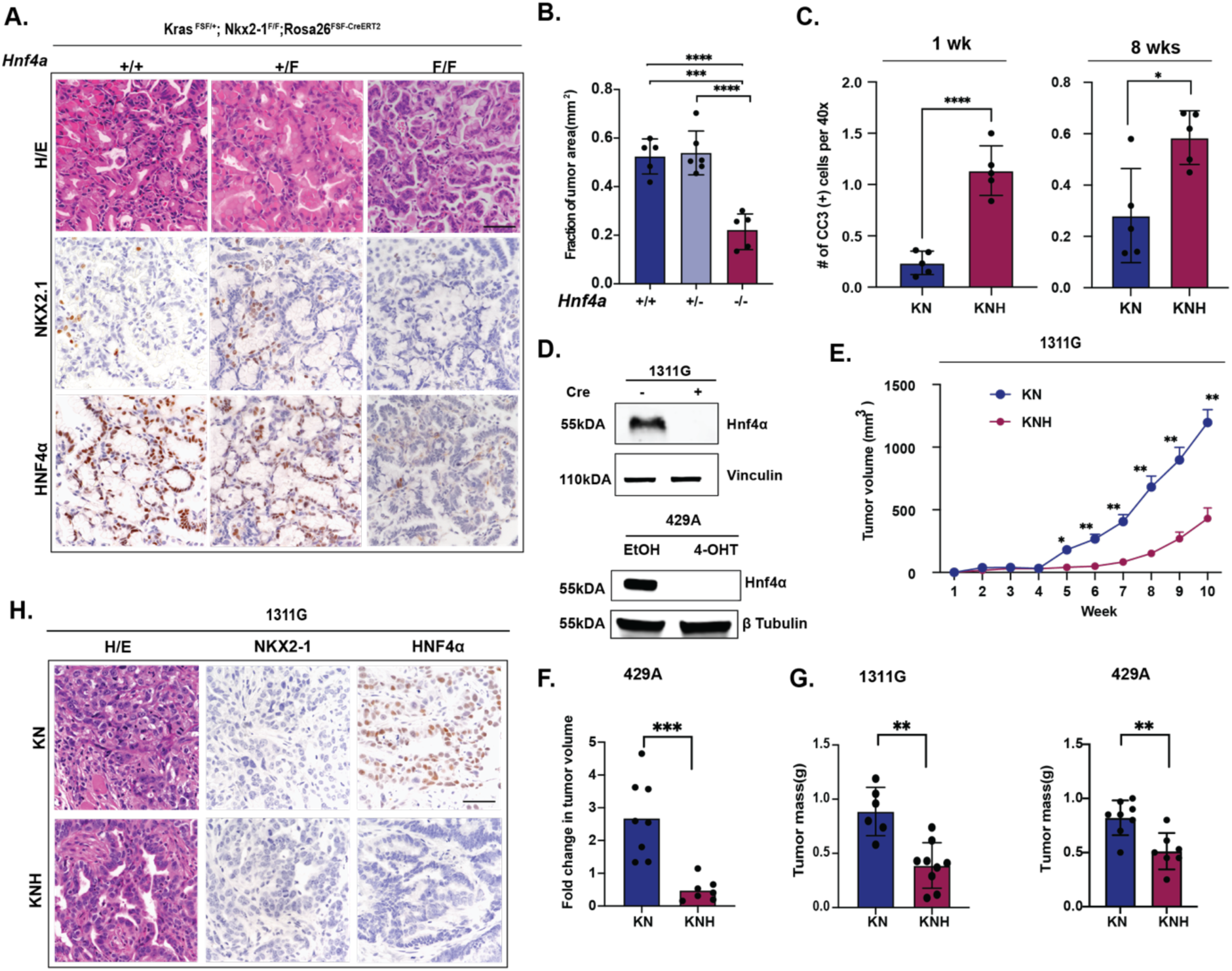
HNF4α is essential for in vivo growth of established IMA. **A.** Representative images of Hematoxylin and Eosin (H&E) staining and immunohistochemistry (IHC) for NKX2-1 and HNF4α in tumors from IMA GEMMs expressing *Hnf4a*^+/+^, *Hnf4a*^+/-^ and *Hnf4a*^-/-^. Scale bar: 100μM. **B.** Quantification of tumor burden in *Hnf4a*^+/+^ (N=5), *Hnf4a*^+/-^ (N=6) and *Hnf4a*^-/-^ (N=5) IMA GEMM cohorts at 14 weeks post-tumor initiation with Ad5mSPC-FlpO (1×10^8^ pfu/mouse). One-way ANOVA showed significant differences among the groups (****p value = <0.0001). Tukey’s multiple comparison test showed: **** p value = <0.0001, *** p value = 0.0001). **C.** Quantification of cleaved caspase-3 (CC3) IHC staining of tumors collected from KN and KNH GEMMs one week (left) or eight weeks (right) after tamoxifen administration. (unpaired t-tests; ****p < 0.0001 and *p = 0.0123). **D.** Representative immunoblots for indicated proteins of 1311G organoids +/- Ad5CMVCre for 3 days (top) or 429A organoids treated with ethanol (EtOH) or 4-hydroxytamoxifen (4-OHT) for 48 hours (bottom) to stably delete *Hnf4a* in vitro. **E.** Longitudinal subcutaneous tumor volume of KN and KNH 1311G organoid lines in NSG mice (unpaired t-tests; *p = 0.0129, **p = 0.0031). **F.** Quantification of fold difference in longitudinal 429A subcutaneous tumor volume in NSG mice before and after 10 days of tamoxifen treatment. Data are presented as the fold difference in tumor volume, comparing pre- and post-treatment measurements (unpaired t-test; ***p = 0.0004). **G.** Quantification of endpoint tumor mass of KN and KNH tumors harvested from NSG mice transplanted with 1311G (left, unpaired t-tests; **p = 0.0013) or 429A (right, unpaired t-tests; **p = 0.0030). **H.** Representative images of H&E staining and IHC for NKX2-1 and HNF4α of subcutaneous tumors derived from allograft transplantation of isogenic 1311G organoid lines with or without HNF4α expression into NSG mice. Scale bar: 100μm.

To uncouple *Hnf4a* deletion from *Nkx2-1* deletion, we derived two isogenic organoid lines from KNH mice (1311G and 429A) that were NKX2-1-deficient and HNF4α-proficient. These lines were subcloned and screened for stochastic retention of HNF4α expression using established protocols(Orstad et al., 2022; Pleguezuelos-Manzano et al., 2020) (**Fig. S1F and Methods**). We then deleted *Hnf4a* in 1311G and 429A lines using Ad5CMV-Cre and 4-hydroxy-tamoxifen (4-OHT), respectively (**Fig**. **1D**). *Hnf4a* deletion did not affect the proliferation of either organoid line in vitro (**Sup. Fig. S1G-J**), but it significantly impaired the growth of both organoid lines in vivo (**Fig. 1E-G**). Histological analysis confirmed that tumors derived from both organoid lines retained their expected morphology and protein expression (**Fig. 1H, Sup. Fig. S1K**). These findings show that HNF4α promotes IMA growth and suppresses apoptosis in vivo.

### HNF4α directly activates a gastric differentiation program in IMA

To understand how HNF4α governs epithelial identity in IMA, we systematically mapped its direct targets and downstream transcriptional programs across multiple models. To enable high-purity isolation of tumor nuclei for chromatin profiling, we incorporated a *Sun1*-*sfGFP* Cre reporter allele(Bhattacharyya et al., 2019; Mo et al., 2015) into KN and KNH IMA GEMMs, allowing efficient isolation of GFP-positive nuclei or whole cells by fluorescence-activated cell sorting (FACS)(Gillis et al., 2023).

Chromatin immunoprecipitation followed by sequencing (ChIP-seq) for HNF4α in GFP-positive nuclei sorted from KN tumors identified 4,021 high-confidence binding sites across two biological replicates (**Supplemental Table S1**). These peaks were predominantly located within intronic, intergenic, and promoter regions (**Fig. 2A**). Motif analysis of these binding sites revealed expected enrichment for HNF4, as well as motifs bound by transcription factor families that regulate endodermal differentiation (e.g. ONECUT, ESRRA and FOX) (**Fig. 2B**). FoxA1/2 regulate gastrointestinal differentiation(Garrison et al., 2006; Kaestner, 2010), and the ONECUT family is crucial for embryonic development of liver, pancreas and neurons(Kropp & Gannon, 2016). ERRα (encoded by *ESRRA*) modulates metabolic processes in GI tissues in cooperation with HNF4α (Cerutti et al., 2023; Scholtes et al., 2024). Furthermore, ENRICHR analysis(E. Y. Chen et al., 2013; Kuleshov et al., 2016) using ARCHS4 tissue annotations demonstrated that HNF4α-bound genes were enriched in signatures of gastrointestinal tissues (**Fig. 2C**).

**Figure 2.**
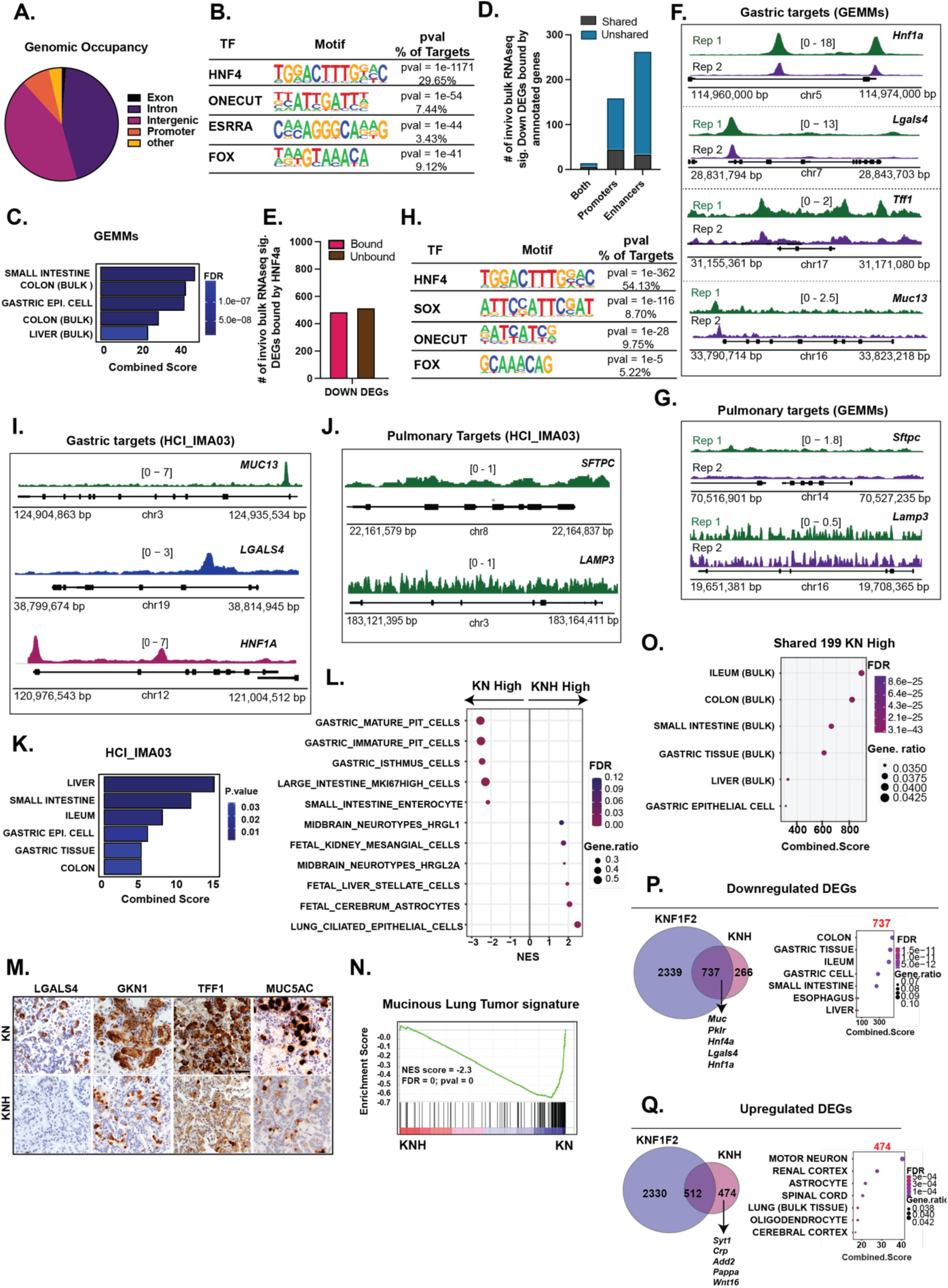
HNF4α directly activates a gastric differentiation program in IMA. **A.** Genome-wide HNF4α ChIP-Seq occupancy in KN GEMMs. **B.** HOMER motif enrichment of HNF4α-bound peaks, showing top transcription factors identified in vivo. **C.** ENRICHR ARCHS4 tissue enrichment of genes annotated from HNF4α peaks in KN. **D.** Bar graph showing the intersection of DEGs from bulk RNA-seq with annotated genes from HNF4α peaks. **E.** Bar plot showing overlap between enhancer, promoter, and dual-mapped HNF4α targets and in vivo downregulated DEGs in the IMA GEMM. **F.** IGV tracks showing HNF4α occupancy at representative gastric lineage genes in KN GEMMs. **G.** IGV tracks showing lack of HNF4α occupancy at representative pulmonary lineage genes in KN GEMMs. **H.** HOMER motif enrichment of HNF4α-bound peaks in HCI_IMA03. **I.** IGV tracks showing HNF4α occupancy at representative gastric lineage genes in HCI_IMA03. **J.** IGV tracks showing lack of HNF4α occupancy at pulmonary lineage genes in HCI_IMA03. **K.** ENRICHR ARCHS4 tissue enrichment of genes annotated from HNF4α peaks in HCI_IMA03. **L.** C8 cell type enrichment analysis using GSEA on DEGs from in vivo KN and KNH tumors. **M.** Representative IHC images of pit cell differentiation markers (LGALS4, GKN1, TFF1, and MUC5AC) in KN and KNH tumors at 14 weeks post-tumor initiation. Scale bar: 100 μm. **N.** GSEA of the mucinous tumor signature of IMA (PMID: 28255028) based on DEGs from bulk RNA-seq of GEMM (KN and KNH) tumors. Normalized enrichment score (NES) and false discovery rate (FDR) are indicated. **O.** ENRICHR ARCHS4 analysis of 199 genes shared between KN-high DEGs and HNF4A-induced genes in H2122 cells reveals enrichment for gastric-related lineages. **P-Q**. Venn diagrams showing overlap of down- and upregulated DEGs between KNF1F2 and KNH tumors. **Q**. Downregulated DEGs showing 737 shared genes (KNF1F2 n = 3,076; KNH n = 1,003; hypergeometric test, p < 1 × 10⁻¹⁵; ∼5-fold enrichment), with ENRICHR ARCHS4 tissue enrichment of shared genes. **R**. Upregulated DEGs showing 512 shared genes (KNF1F2 n = 2,842; KNH n = 986; p < 1 × 10⁻¹⁹⁴; ∼3.8-fold enrichment), and 474 genes uniquely upregulated in KNH with ENRICHR ARCHS4 enrichment.

To assess the functional consequences of HNF4α binding, we performed bulk RNA-seq on GFP-positive tumor cells isolated from KN and KNH GEMMs (n = 4 mice per genotype), identifying 1,903 differentially expressed genes (DEGs; log₂FC > 0.585, padj < 0.05; **Supplemental Table S2**). Integration of ChIP-seq and RNA-seq revealed that ∼50% of genes downregulated and ∼15% of genes upregulated in KNH tumors were direct HNF4α targets (**Fig. 2D**), consistent with HNF4α functioning predominantly as a direct transcriptional activator in IMA. To more precisely associate HNF4a binding sites with target genes, we intersected these data with H3K27ac HiChIP data from in vivo KN tumors(Gillis et al., 2023). Approximately 50% of HNF4α-bound regions that looped to a promoter, enhancer, or both were linked to genes that were downregulated upon *Hnf4a* deletion in vivo, as determined by bulk RNA-seq (**Fig. 2E, Supplemental Table S3**), reinforcing a model in which HNF4α sustains lineage-specific transcription through enhancer activation.

Inspection of individual genes from in vivo KN tumors showed strong HNF4α binding at regulatory elements of canonical gastric genes including *Hnf1a, Lgals4, Tff1* and *Muc13* (**Fig. 2F-G**). Importantly, HNF4α ChIP-seq in an IMA-derived organoid (HCI_IMA03) demonstrated similar motif enrichment, target gene occupancy, and enrichment for gastrointestinal tissue signatures (**Fig. 2H-K, Sup. Fig. S2A-C**). Similar HNF4α binding profiles were also observed in two mouse IMA organoid lines (429A and 1311G) (**Sup. Fig. S2D-L**, **Supplemental Table S1**). Taken together, these data demonstrate that HNF4α chromatin binding patterns in murine IMA models recapitulate the human disease. We did not observe significant HNF4α binding at pulmonary marker genes in murine or human IMA models. This gastric-restricted binding pattern contrasts with the hybrid-identity LUAD cell state, in which we previously found that HNF4α can co-localize with NKX2-1 at pulmonary gene loci(Fort et al., 2024).

To identify specific HNF4α-regulated identity programs in IMA, we performed Gene Set Enrichment Analysis (GSEA)(Liberzon et al., 2011; Subramanian et al., 2005) of DEGs against cell type signatures. This analysis revealed that gene signatures associated with gastric cell types, including immature and mature pit cells as well as isthmus cells, were markedly depleted in KNH tumors compared to KN tumors, whereas non-gastric identity programs, including astrocyte, neuronal, and liver signatures, were upregulated in KNH (**Fig. 2L, Sup. Fig. S3A; Supplemental Table S4**). IHC validated a reduction in the pan-gastric marker galectin-4 (LGALS4) as well as the pit cell markers gastrokine-1 (GKN1), trefoil factor 1 (TFF1) and MUC5AC in tumors lacking HNF4α (**Fig. 2M**). Additionally, *Hnf4a* deletion caused a significant decline in an IMA gene expression signature derived from human tumors(Guo et al., 2017) (**Fig. 2N**). We also compared KN-high DEGs (defined as genes expressed at significantly higher levels in KN tumors relative to KNH; log₂FC > 0.585, padj < 0.05, **Supplemental Table S2**) with genes induced by exogenous HNF4α in the NKX2-1/HNF4α dual-negative H2122 human lung carcinoma cells(Stuart et al., 2024). We identified 199 shared genes, which were enriched for gastric-related cell lineages when compared to the ARCHS4 resource via ENRICHR analysis (**Fig. 2O**). Taken together, these data show that DEGS identified in GEMMs are relevant to human IMA.

Bulk RNA-seq of KN and KNH organoids demonstrated significant overlap with the in vivo GEMM. Bulk RNA-seq of 429A and 1311G organoids revealed 818 DEGs and 2,323, respectively, between KN and KNH genotypes (log₂FC > 0.585, padj < 0.05, **Supplemental Tables S5-S6**). The 429A organoid model showed the highest concordance with tumor cell-derived DEGs, with 107 KNH-high genes (39%) and 202 KN-high genes (60%) overlapping with upregulated genes in KNH and KN tumor cells, respectively. The 1311G organoid line showed more modest overlap, with 131 KNH-high (12%) and 284 KN-high (23%) shared with in vivo profiles (**Sup. Fig. S3B-F**). GSEA of organoid DEGs from both models mirrored in vivo trends, showing depletion of gastric pit cell identity and activation of non-gastric signatures in the KNH tumors (**Sup. Fig. S3G-H, Supplemental Table S7**). Together, these findings demonstrate that HNF4α safeguards gastric lineage fidelity in IMA by enforcing gastric pit cell identity and suppressing alternative identities, as shown across these orthogonal models of IMA.

Our prior work showed that FoxA1/2 are required for *Hnf4a* expression in IMA, suggesting that a FoxA1/2-> HNF4α transcription factor hierarchy controls IMA identity(Gillis et al., 2023). To define the subset of FoxA1/2-dependent genes that are activated by HNF4α, we compared DEGs from KNH tumors to those from *Foxa1/2* double-knockout tumors (KNF1F2)(Gillis et al., 2023), both relative to KN tumors. Among 737 shared downregulated genes, most were enriched in gastrointestinal pathways (**Fig. 2P**), and this overlap was highly significant by hypergeometric test (*p* < 1×10⁻¹⁵; ∼5-fold enrichment)(Johnson et al., 2005). These data show that HNF4α is a major downstream effector of FoxA1/2 for gastric identity specification in IMA. Intriguingly, we identified 474 genes uniquely upregulated in KNH but not in KNF1F2, many of which were associated with non-GI lineages, suggesting FoxA1/2 are required to activate non-GI identity programs in KNH tumors (**Fig. 2Q**).

Finally, pathway analysis of in vivo tumor cell-derived DEGs showed that HNF4α loss also modulates metabolic gene programs. KN tumors were enriched for xenobiotic metabolism, steroid biosynthesis, glycolysis, and G2-M checkpoint pathways (KEGG, Hallmark, **Sup. Fig. S3I-J, Supplemental Table S8**) while KNH tumors showed enrichment for cilium assembly and organelle biogenesis (Reactome, **Sup. Fig. S3K, Supplemental Table S8**). These results were consistent with gene ontology findings from the organoid models (**Supplemental Table S9**) and reflect canonical roles of HNF4α as a key regulator of multiple metabolic programs, including xenobiotic metabolism, in the GI tract and liver(Deans et al., 2023; Hwang-Verslues & Sladek, 2010; Thymiakou et al., 2023). Together, these data support HNF4α as a master regulator of gastric identity in IMA, acting through enhancer-mediated activation of pit cell genes and repression of transcriptional plasticity.

### HNF4α Restricts Cellular Heterogeneity and Prevents Lineage Deviation in IMA

Previous studies have highlighted the contribution of intercellular heterogeneity to lineage plasticity(Han et al., 2024) and therapeutic resistance in LUAD, including IMA(Maleki et al., 2024; Zewdu et al., 2021). Although bulk RNA-seq revealed that *Hnf4a* deletion broadly suppresses gastric identity and activates non-gastric programs, this approach averages across cell states and obscures the spectrum of transcriptional heterogeneity present within tumors.

To investigate the role of HNF4α in distinct IMA subpopulations and assess whether its deletion alters the cellular composition or differentiation trajectory of IMA, we performed single-cell RNA sequencing (scRNA-seq) on sorted GFP-positive tumor cells from KN and KNH tumor cells 14 weeks after tumor initiation (n = 2 mice per genotype, **Sup. Fig. S4A-D**). Analysis of all high-quality cells demonstrated that most were bona fide tumor cells, defined by co-expression of CreERT2 and nuclear-localized sfGFP (**Sup. Fig. S4E-F**). After excluding the small number of normal cells (sfGFP/CreERT2-negative), we identified 18 clusters comprising 5,003 KN cells and 5,024 KNH tumor cells (**Fig. 3A; Supplemental Table S10**). Louvain clustering further resolved these tumor cells into two major transcriptional groups: Group A, with roughly equal contribution from KN and KNH cells, and Group B, which comprised ∼80% of tumor cells and had a slightly higher proportion of KN cells (**Fig. 3B-E**). Notably, unrecombined tumor cells (cluster 12) constituted less than 2% of the dataset and expressed both alveolar type 2 (AT2) gene signature and markers such as *Sftpc*, *Sftpa and Nkx2-1*(Marjanovic et al., 2020).(**Sup. Fig. S4G-H**). Quantification of exon-specific reads confirmed efficient *Hnf4a* deletion in KNH cells (**Sup. Fig. S4I; Supplemental Table S10**).

**Figure 3.**
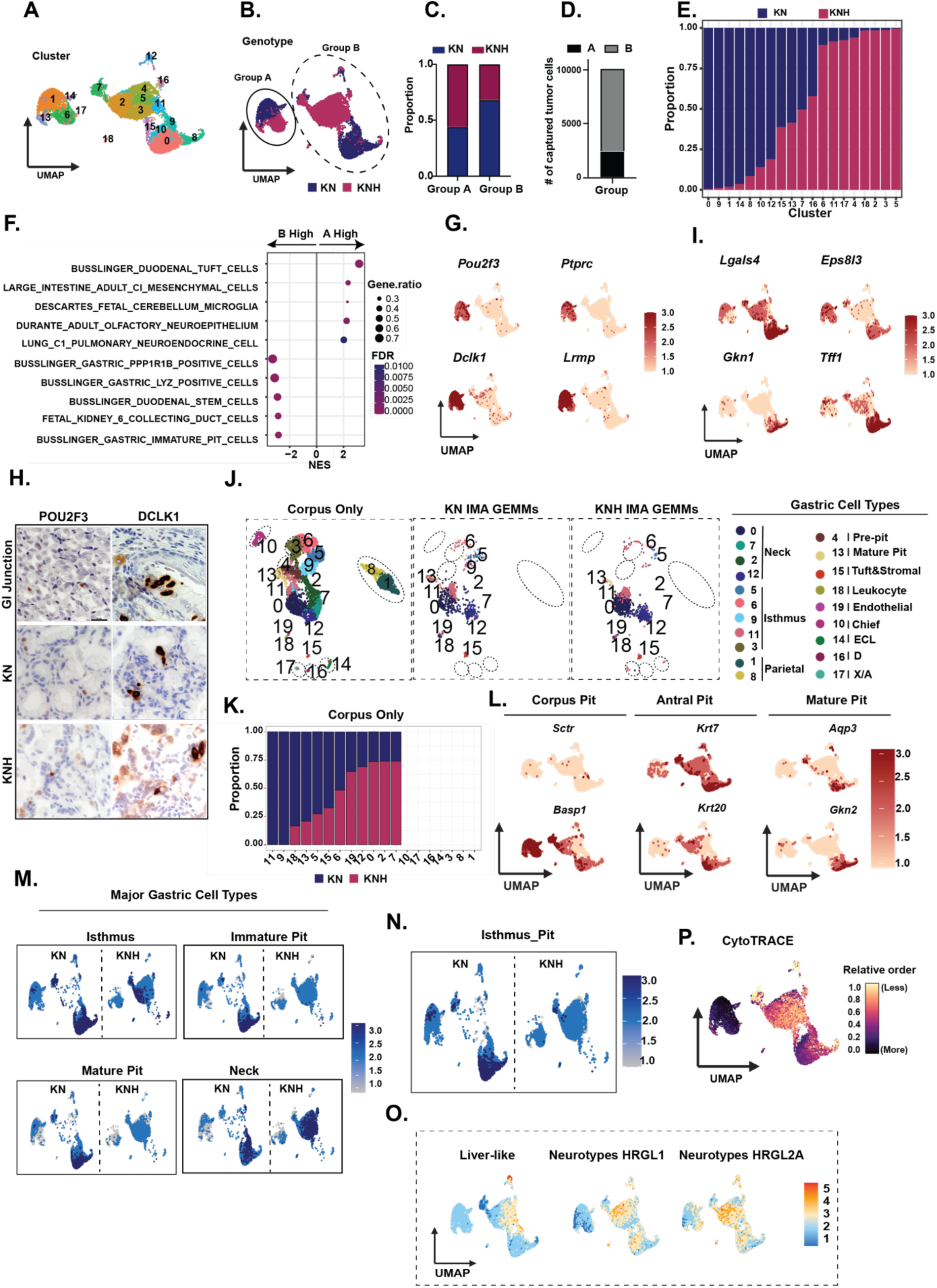
HNF4α Restricts Cellular Heterogeneity and Prevents Lineage Deviation in IMA. **A.** UMAP of KN (5,001 cells) and KNH (5,024 cells) tumors (n = 2 mice per genotype, multiple tumors per mouse) colored by Seurat-defined clusters. **B.** UMAP of KN and KNH tumor cells colored by genotype. **C.** Proportion of KN and KNH tumor cells captured in group A and B. **D.** Total number of tumor cells captured in group A and B. **E.** Proportion of bona fide tumor cells from each genotype across Seurat-defined clusters. **F.** C8 cell type enrichment analysis of DEGs comparing Group A to Group B **G.** UMAPs showing relative expression of representative tuft cell marker genes (*Pou2f3*, *Ptprc, Dclk1* and *Lrmp*) across tumor cells. **H.** Representative IHC staining for DCLK1, POU2F3, and PTPRC in KN and KNH tumors at 14 weeks post-tumor initiation. Scale bar: 100 μm **I.** UMAPs showing relative expression of representative HNF4α target genes (*Lgals4, Eps813, Gkn1*, and *Tff1*) across tumor cells. **J.** UMAP of Takada et al. (PMID: 37386010) corpus only dataset annotated for gastric epithelial and stromal cell types (left), alongside KN and KNH tumor cells mapped to the same reference using Seurat label transfer (right). **K.** Proportion of KN and KNH tumor cells mapping to each cluster in corpus only dataset above. **L.** Feature plots showing expression of corpus pit, antral pit, and mature pit cell markers in IMA tumor cells. **M.** UMAPs of gene module scores for major gastric cell types, including isthmus, immature pit, mature pit and neck cells. **N.** UMAPs of gene module scores for cells undergoing differentiation along the isthmus-to-pit trajectory as described by Takada et al. (PMID: 37386010). **O.** UMAPs showing enrichment of gene modules associated with alternative cell types upregulated in KNH tumors. **P.** UMAP depicting the relative differentiation potential of KN and KNH tumor cells using the CytoTRACE algorithm.

Differential expression analysis identified 2,958 DEGs between Groups A and B (1,580 Group A high, and 1,378 Group B high; log₂FC > 0.25; **Supplemental Table S10**). Group A was enriched for a tuft cell program, marked by the master regulator *Pou2f3* as well as other tuft cell markers such as *Dclk1, Ptprc (CD45)*, and *Lrmp* (**Fig. 3F-G**). These tuft markers were expressed at comparable levels between KN and KNH tumors based on scRNA-seq and IHC, which confirmed POU2F3 and DCLK1 protein expression in a minority of tumor cells of both genotypes (**Fig. 3H**). Moreover, *Hnf4a* deletion did not affect the expression of either neuronal or immune-associated markers that have been associated with specific subpopulations of normal tuft cells (**Sup. Fig. 3J**). These data show that the KN and KNH IMA models harbor a stable tuft-like subpopulation, which we have previously observed in human IMA and BRAF^V600E^-driven GEMMs(Zewdu et al., 2021).

Despite similar expression of major tuft markers, KN and KNH cells within Group A clustered distinctly. We identified 975 DEGs between KN (n = 1,631) and KNH (n = 761) cells in Group A (log₂FC > 0.25; **Supplemental Table S10**). Excluding genes also differentially expressed in Group B, we identified 467 KN-high and 281 KNH-high genes unique to Group A (**Sup. Fig. S4K**). Surprisingly, cell type enrichment analysis of these unique DEGs revealed reduced signatures for tuft cells, microglia, and neuronal lineages in KNH cells (**Sup. Fig. S4L**). Leading edge analysis via GSEA showed that the loss of tuft cell signature in Group A was driven by reduced expression of a subset of tuft-associated genes including IL17rb(Goto et al., 2019), Hpgds(Xiong et al., 2022), Alox5(McGinty et al., 2020) and Plcg2(Xiong et al., 2022) (**Sup. Fig. S4M**). ENRICHR analysis confirmed that these genes are functionally linked to tuft cell differentiation, underscoring the biological significance of this transcriptional shift (**Sup. Fig. S4N**). These data show that HNF4α regulates a distinct set of genes in tuft-like IMA cells, including a subset of normal tuft cell markers, suggesting that loss of HNF4α might impair the functional properties of these cells without fundamentally altering their identity.

We next evaluated Group B, which was enriched for gene signatures associated with gastric pit cell differentiation. HNF4α targets identified in bulk RNA-seq analysis, including gastric markers *Lgals4*, *Eps8l3*, *Gkn1* and *Tff1,* were significantly downregulated in Group B KNH cells (**Fig. 3I**). To evaluate the extent to which IMA tumor cells recapitulate normal gastric differentiation, we projected scRNA-seq profiles onto gastric epithelial reference atlases(Takada et al., 2023). In the corpus-only atlas, KN cells mapped primarily to mature pit (cluster 13), neck (clusters 0, 2, 7, and 12), and isthmus (clusters 5, 6, 9, and 11) cell types, with minor representation of tuft cells (cluster 15) (**Fig. 3J-K**). There was a notable absence of KN cells mapping to normal parietal, chief and enteroendocrine cells. In contrast, KNH cells were underrepresented in pit and isthmus clusters and overrepresented in neck cell clusters. Similar differences were observed when tumor cells were mapped to a second atlas that includes both corpus and antrum (**Sup. Fig. S5A-B**).

Analysis of specific marker genes as well as cell type signatures(Busslinger et al., 2021) demonstrated a clear loss of pit and isthmus identity in Group B KNH cells (**Fig. 3L-M, Sup. Fig. S5C**). We further examined differentiation dynamics using established isthmus-to-pit transition markers(Takada et al., 2023), which showed that markers of progression into more differentiated pit cells were significantly diminished, indicating that HNF4α is required for this lineage commitment step(Takada et al., 2023; Willet & Mills, 2016) (**Fig. 3N**). Although a higher proportion of KNH cells mapped to neck cell clusters, the level of the neck cell signature was similar between genotypes. This may be due to the fact that some neck markers, such as *Tff2*, are expressed at high levels in both neck and pit cells (**Sup. Fig. S5D**) and suggest that KNH cells exhibit predominantly a loss of pit/isthmus identity rather than an increase in neck cell identity.

Finally, we asked whether the non-GI identities observed in KNH cells by bulk RNA-seq data were detectable at the single cell level. Group B KNH tumors exhibited specific enrichment of curated non-GI signatures, including neuronal and liver-like programs (**Fig. 3O**). These signatures encompassed both canonical marker genes (e.g., *Fgfr3, Kcnh7, Rora* for neuronal; *Mgst1, Trf, Cp* for liver) and broader transcriptional networks associated with these cell types (**Sup. Fig. S5E**). Moreover, Group B KNH cells exhibited less differentiated, more progenitor-like state than Group B KN cells by CytoTRACE analysis(Gulati et al., 2020), a computational metric that infers cellular differentiation potential based on transcriptional diversity (**Fig. 3P**). In contrast, group A cells of both genotypes were predicted to be in a highly differentiated state based on this analysis. This suggests that HNF4α loss promotes dedifferentiation specifically in pit/isthmus-like IMA cells, but not in tuft-like IMA cells.

To test whether this dedifferentiation reflected reactivation of fetal or pluripotent transcriptional programs in KNH, we analyzed expression of iPSC/ESC gene signatures(Mizuno et al., 2010), canonical fetal intestinal stem cell markers (*Ly6a*, *Tacstd2*, *Anxa1*, *Flt1*)(Yui et al., 2018), and the MSigDB DESCARTES fetal intestine epithelial gene sets(Cao et al., 2020). All were similarly expressed in KN and KNH cells in Group B, indicating that the dedifferentiated state in KNH likely does not reflect activation of a fetal-like or pluripotent program (data not shown).

Taken together, these data show that HNF4α regulates the transcriptome of both major (pit/isthmus-like) and minor (tuft-like) IMA subpopulations, with both overlapping and unique targets in each subpopulation. In the major subpopulation, *Hnf4a* deletion leads to dedifferentiation marked by a loss of pit-like identity and induction of non-GI signatures. In contrast, the identity of the tuft-like subpopulation is more stable upon *Hnf4a* deletion, although a subset of tuft cell-associated transcripts exhibits HNF4α dependency.

### HNF4α Loss Reprograms FoxA1/2 Binding to Drive Non-Gastric States in IMA

Having established that HNF4α is required for IMA growth (**Fig. 1**), directly activates gastric differentiation programs (**Fig. 2**), and restricts transcriptional plasticity at the single-cell level (**Fig. 3**), we next investigated how its loss reconfigures the underlying transcriptional network. scRNA-seq analyses revealed that *Hnf4a* deletion promotes the emergence of non-gastric lineage programs, including neuronal- and liver-like states, raising the question of which transcription factors drive these alternate identities.

FoxA1/2 are diffusely expressed in both KN and KNH tumors (**Sup. Fig. S6A**), and FOXA motifs were highly enriched at HNF4α-bound regulatory regions (**Fig. 2B, 2H, Sup. Fig. S2E, S2I**), suggesting potential cooperation or hierarchical regulation. Bulk RNA-seq across multiple IMA models revealed that many of the gene expression changes observed upon *Hnf4a* loss including activation of non-gastric programs, were partially dependent on FoxA1/2 activity (**Fig. 2Q**). These findings led us to hypothesize that in the absence of HNF4α, FoxA1/2 engage de novo chromatin sites to pioneer non-GI programs and rewire cell state identity.

To test this, we performed ChIP-seq for FoxA1 and FoxA2 using GFP-positive nuclei sorted from lung tumors of KN and KNH GEMMs. HOMER motif analysis revealed that a FOXA binding motif was enriched in peaks from both KN and KNH tumors, whereas HNF4α motifs were only enriched in KN peaks (**Sup. Fig. S6B, Supplemental Table S1**), consistent with co-occupancy at gastric regulatory loci in KN tumors. Similar trends were observed for FoxA2 (**Sup. Fig. S6C, Supplemental Table S1**), with extensive FoxA1/2 co-occupancy across both genotypes (**Sup. Fig. S6D-F**). These co-binding events were exemplified at the *Hnf1a* locus, where IGV visualization revealed overlapping FoxA1, FoxA2, and HNF4α binding in KN tumors (**Sup. Fig. S6G**). FoxA1 ChIP-seq in 1311G KN and KNH organoids recapitulated these in vivo patterns (**Sup. Fig. S6H-K**).

Using DiffBind, we identified 2,369 KN-specific and 494 KNH-specific in vivo FoxA1 peaks, and 1,443 KN-specific and 983 KNH-specific in vivo FoxA2 peaks (**Supplemental Table S11**). Mapping these differential FoxA1 peaks onto the merged binding sets for HNF4α (in KN) and FoxA1/2 (in KN and KNH) revealed striking patterns: KN-specific FoxA1 peaks overlapped substantially with FoxA1, FoxA2, and HNF4α sites in KN, while KNH-specific peaks overlapped with FoxA1 and FoxA2 in KNH but did not overlap with HNF4α (**Fig. 4A**). Similar observations were made for FoxA2 (**Sup. Fig. S7A, Table S11**).

**Figure 4.**
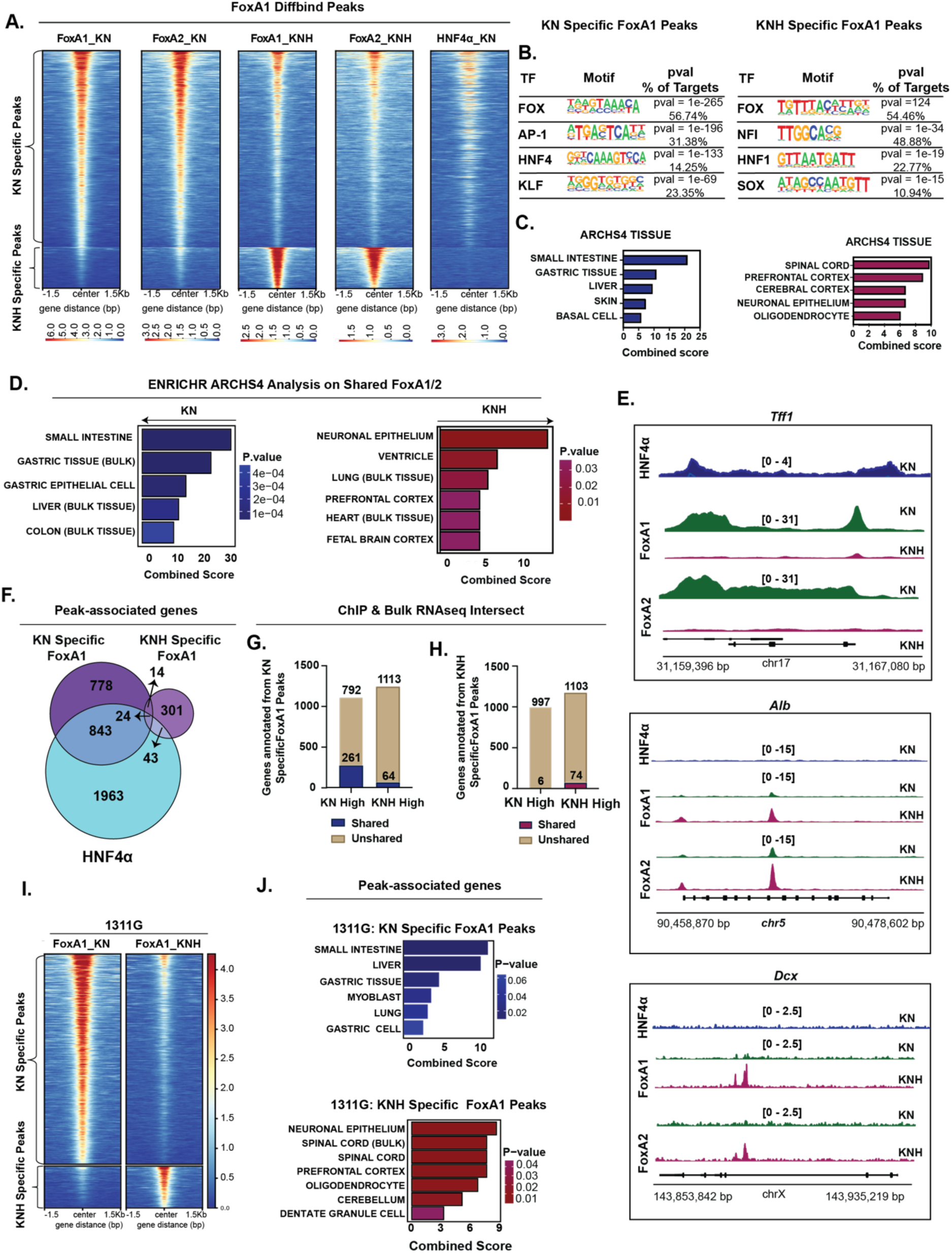
HNF4α Loss Reprograms FoxA1/2 Binding to Drive Non-Gastric States in IMA. **A.** Heatmap showing differential FoxA1 signal enrichment between KN and KNH tumors in vivo, identified using DiffBind (adjusted p < 0.05). Differential signal intensities were quantified over merged peak regions, defined as significant peaks (MACS2, adjusted p < 0.05) detected in at least one condition, spanning HNF4α, FoxA1, and FoxA2 in KN, and FoxA1 and FoxA2 in KNH. **B.** HOMER motif analysis of transcription factor motifs enriched at differential FoxA1 peaks. Left: KN-specific motifs. Right: KNH-specific motifs. **C.** ENRICHR ARCHS4 tissue enrichment of genes annotated from differential FoxA1 peaks. Left: KN-specific. Right: KNH-specific. **D.** ENRICHR ARCHS4 tissue enrichment of genes annotated from differential FoxA1/2 peaks. Left: KN-specific. Right: KNH-specific. **E.** IGV tracks showing examples of differential FoxA1/2 occupancy in KN and KNH tumors. Binding is lost at the gastric gene *Tff1* and gained at the neuronal gene *Dcx* and liver gene *Alb* in KNH tumors. **F.** Venn diagram showing the overlap between genes associated with differential FoxA1 peaks and those linked to HNF4α-bound regions in KN tumors in vivo. **G.** Bar plot showing overlap between genes annotated from KN-specific FoxA1 peaks and DEGs from bulk RNA-seq. **H.** Bar plot showing overlap between genes annotated from KNH-specific FoxA1 peaks and DEGs from bulk RNA-seq in vivo. **I.** Heatmap showing FoxA1 occupancy at differential binding sites in KN versus KNH following HNF4α loss in 1311G organoids. DiffBind analysis identified 1,799 gained (Cluster 1, KN-specific) and 337 lost (Cluster 2, KNH-specific) peaks. **J.** ENRICHR ARCHS4 tissue enrichment of genes annotated from differential FoxA1 peaks in vitro. Top: 1311G KN-specific peaks. Bottom: 1311G KNH-specific peaks.

HOMER motif analysis revealed that KN-specific FoxA1 peaks were enriched for motifs corresponding to FOX, AP-1, HNF4, and KLF transcription factor families, while KNH-specific peaks were enriched for FOX, NFI, HNF1, and SOX motifs (**Fig. 4B**). The presence of HNF4, AP-1 and KLF motifs in KN-specific peaks suggests that FoxA1 cooperates with these factors to drive gastric identity in IMA(Li et al., 2023; Moore et al., 2016; Wang et al., 2017). In contrast, the enrichment of NFI and SOX motifs in KNH-specific peaks suggests that, in the absence of HNF4α, FoxA1/2 may cooperate with alternative transcriptional partners to drive lineage plasticity and activate non-gastric programs(Fu & Shi, 2017; Malaymar Pinar et al., 2025; Mason et al., 2009). FoxA2 motif analysis showed a similar pattern (**Sup. Fig. S7B**).

ENRICHR cell type enrichment analysis on genes annotated with KN- and KNH-specific FoxA1/2 peaks showed that genes bound by KN-specific FoxA1/2 peaks were enriched for gastric epithelial lineages, whereas those bound by KNH-specific FoxA1/2 peaks were linked to neuronal and hepatic cell types (**Fig. 4C, Sup. Fig. S7C**). Similar results were obtained when analyzing genotype-specific shared FoxA1/2 binding sites (**Fig. 4D**). For example, FoxA1/2 binding at *Tff1* was lost in KNH, while binding increased at representative neuronal (*Dcx)* and hepatic (*Alb)* marker genes (**Fig. 4E**). These binding changes were accompanied by corresponding shifts in gene expression: genes associated with KN-specific FoxA1 peaks were significantly downregulated in KNH tumors, while those linked to KNH-specific FoxA1 peaks were upregulated (**Fig. 4F-H**). Similar transcriptional correlations were observed for FoxA2 (**Sup. Fig. S7D-F**).

DiffBind differential binding analysis of FoxA1 ChIP-seq in KN and KNH 1311G organoids revealed substantial relocalization from canonical gastric loci, including HNF4α target genes, in KN organoids to non-gastric loci associated with neuronal cell types in KNH organoids (**Sup. Fig. S7G-H, Fig. 4I-J**). These shifts in FoxA1 occupancy mirrored the enrichment of non-gastric gene cell types observed in KNH across both scRNA-seq and bulk RNA-seq datasets (**Fig. 3O & Fig. 2L**).

These data support a model in which HNF4α promotes FoxA1/2 binding to gastric regulatory elements in IMA, and HNF4α loss permits FoxA1/2 redistribution to the regulatory elements of genes associated with alternative identities, providing a molecular mechanism for the dedifferentiation of KNH tumor cells. The extensive overlap between FoxA1 and FoxA2 peaks in both genotypes further underscore their redundant and cooperative roles in the regulation of IMA identity, even in the HNF4α -negative state.

### HNF4α dampens response of IMA to KRAS inhibition

Drug-tolerant persister (DTP) cells(He et al., 2024) represent a major barrier to durable responses in oncogene-driven cancers, including LUAD(Moghal et al., 2023), where they facilitate survival under targeted therapy pressure. Recent work has shown that gastrointestinal and mucinous gene programs are enriched in DTP states following RAS inhibitor treatment(Araujo et al., 2024), suggesting a potential link to gastric lineage regulators. To assess whether HNF4α influences therapeutic response in IMA, we examined the expression of gastrointestinal and mucinous genes previously linked to RAS inhibitor-induced DTP states in LUAD. Approximately 80% of these genes were downregulated in KNH tumors, suggesting that HNF4α maintains transcriptional programs associated with the DTP phenotype (**Fig. 5A**). These findings prompted us to test whether HNF4α directly modulates the response of IMA to KRAS^G12D^ inhibition.

**Figure 5.**
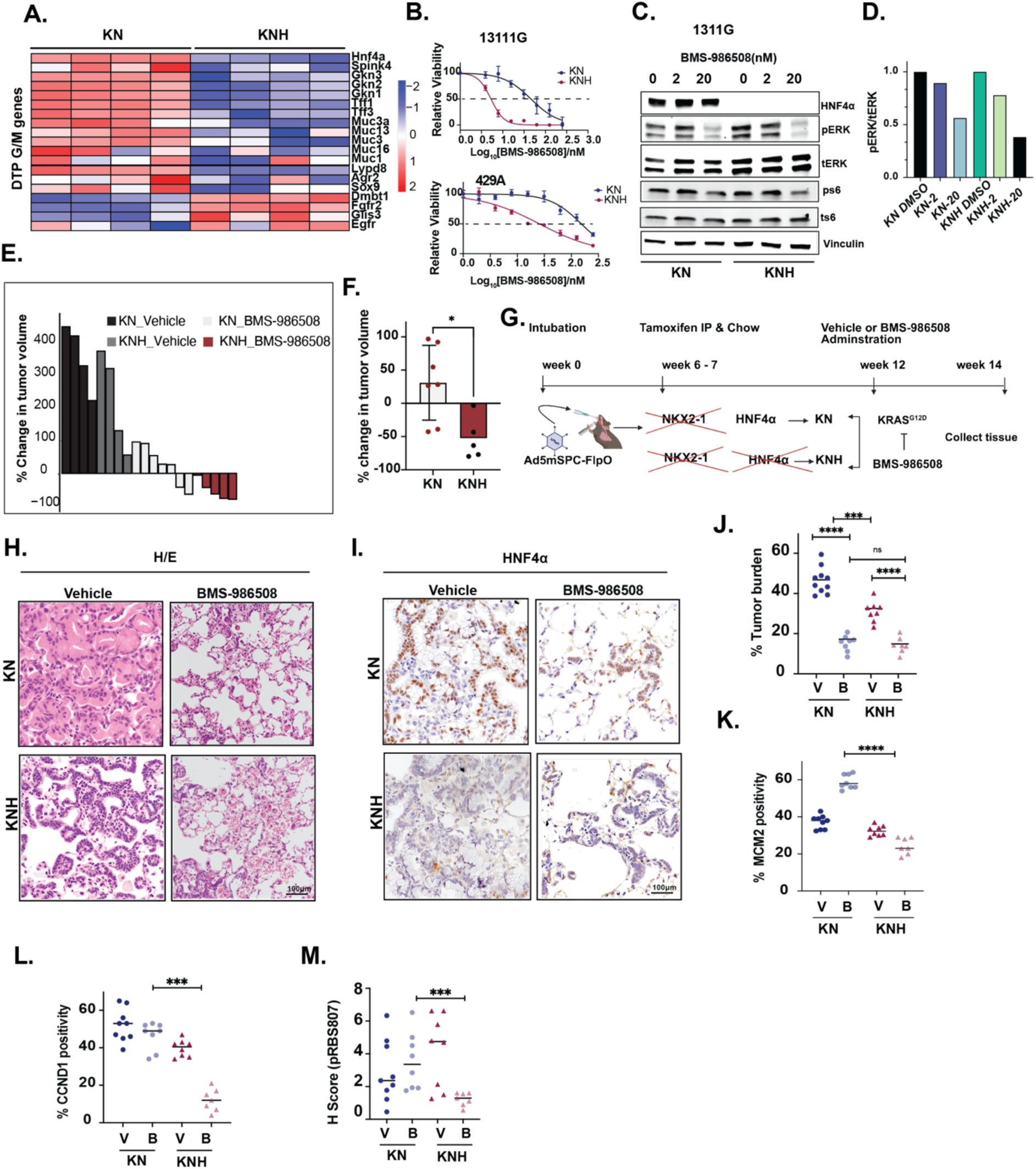
HNF4α dampens response of IMA to KRAS inhibition. **A.** Heatmap showing log₂-normalized expression of GI/mucinous genes that persist in drug-tolerant persister (DTP) cells (PMID: 38975897) in KN and KNH tumors in vivo. **B.** Dose-response curves for KN and KNH IMA organoids treated with BMS-986508. Left: 1311G organoids (KN IC_50_= 45.75 nM, KNH IC_50_= 5.27 nM). Right: 429A organoids (KN IC_50_= 152.3 nM, KNH IC_50_= 25.61 nM). Data represent one of three biological replicates; error bars indicate SEM. **C.** Immunoblot of 1311G KN and KNH organoids treated for 2 hours with vehicle or increasing doses of BMS-986508, probed for the indicated proteins. **D.** Quantification of pERK/tERK ratios for each sample in panel B. **E.** Waterfall plot showing fold change in tumor volume for individual NSG mice bearing subcutaneous KN or KNH 1311G tumors. Mice were treated by daily intraperitoneal (i.p.) injection with 10 mg/kg BMS-986508 or vehicle for 7 days, followed by 14 days at 30 mg/kg, then weaned for 14 days. **F.** Comparison of tumor volumes in KN and KNH allografts before and after BMS-986508 treatment. No significant difference at baseline (p = 0.2020); post-treatment, KNH tumors were significantly smaller than KN (p = 0.0480). Median volumes (mm³): KN baseline = 237.2, KNH baseline = 165.9; KN post = 284.7, KNH post = 68.95. **G.** Schematic of in vivo BMS-986508 treatment protocol in GEMMs. **H.** Representative H&E images of KN and KNH tumors from GEMMs treated with vehicle or 30 mg/kg BMS-986508 for 14 days. Scale bar: 100 μm. **I.** Representative IHC images for HNF4α in KN and KNH tumors from GEMMs treated. Scale bar: 100 μm. **J.** Tumor burden quantification in individual KN and KNH GEMM mice following two weeks of treatment with 30 mg/kg BMS-986508 or vehicle (Mann-Whitney test: ns = 0.6943, ***p = 0.0002, ****p < 0.0001). **K-L.** Quantification of MCM2 (**K**) and CCND1 (**L**) expression by IHC in lung tumors from KN and KNH mice treated with BMS-986508 or vehicle. Each data point represents the average percentage of positive tumor cells per mouse (across multiple tumors). Statistical analysis was performed using the Mann–Whitney test (****p < 0.0001 for MCM2; ***p = 0.0003 for CCND1). **M.** H-SCORE quantification of pRBS807 expression via IHC in lung tissues from KN and KNH mice (Mann-Whitney test, ***p = 0.0003). Each data point represents the average percentage of positive tumor cells per mouse (across multiple tumors).

Treatment of 1311G and 429A organoids with the KRAS^G12D^ inhibitor BMS-986508 (formerly as MRTX1133) demonstrated that *Hnf4a* deletion reduced the IC_50_ of BMS-986508 in 1311G by 9-fold and in 429A by 5-fold (**Fig. 5B**). BMS-986508 inhibited KRAS effectors in both genotypes (KN and KNH), although the effect was slightly more pronounced in KNH organoids (**Fig. 5C-D**).

To evaluate in vivo responses, we subcutaneously implanted 1311G KN and KNH cells into NSG mice and treated them with vehicle or BMS-986508. Because KNH tumors grow more slowly than KN tumors under untreated conditions (**Fig. 1E**), we initiated treatment when tumor volumes were equivalent across genotypes to ensure a comparable baseline for assessing drug response (**Sup. Fig. S8A**). Most KNH tumors regressed in response to BMS-986508, whereas 5/7 KN tumors continued to grow, albeit at a slower rate than vehicle controls (**Fig. 5E-F, Sup. Fig. S8B,**). Taken together, these data showed that *Hnf4a* deletion sensitized IMA to BMS-986508 both in vitro and in vivo.

To further assess response in an immune-competent setting, we treated KN and KNH GEMM mice harboring autochthonous tumors with 30 mg/kg BMS-986508 or vehicle for 14 days (**Fig. 5G**). Post-treatment histological analysis revealed that BMS-986508 significantly reduced the overall tumor burden in both genotypes and suppressed pERK levels, confirming effective pathway inhibition. HNF4α expression remained comparable in KN tumors treated with vehicle or BMS-986508, and residual tumors in KNH mice did not exhibit selective enrichment for HNF4α positive cells (**Fig. 5H-J, Sup. Fig. S8C**). While residual tumor burden did not differ significantly between KN and KNH mice at this time point, we observed a striking divergence in cell cycle behavior. Consistent with response to targeted therapy in a BRAF^V600E^-driven IMA model(Zewdu et al., 2021), BMS-986508 induced a paradoxical increase in the percentage of MCM2-positive DTPs and did not reduce Cyclin D1 and phosphorylated Rb (pRb-S807) levels in KN tumors (**Fig. 5K-M, Sup. Fig. S8D-F**). These data indicate that, despite ERK inhibition and tumor regression, DTPs in KN tumors fail to exit the cell cycle. In contrast, residual tumors in KNH mice exhibited significantly lower MCM2, Cyclin D1, and pRB-S807 positivity, showing that HNF4α loss enables a higher proportion of DTPs to exit the cell cycle in response to KRAS inhibition.

Together, these findings support a model in which HNF4α prevents quiescence in IMA DTPs by maintaining Cyclin D1 expression and Rb phosphorylation, despite KRAS/ERK inhibition. Notably, in one BMS-986508-treated KN tumor, we observed histological evidence of transdifferentiation from a mucinous to squamous phenotype, consistent with emerging reports of lineage plasticity under KRAS inhibition(Awad et al., 2021; Tong et al., 2024), although this was an isolated observation (**Sup. Fig. S8G**).

### HNF4α impairs response to KRAS inhibition by maintaining NRF2 activity in IMA

Given that BMS-986508 suppressed pERK in both genotypes, we sought to identify alternative mechanisms underlying differential sensitivity to KRAS inhibition. Pathway analysis revealed that *Hnf4a* deletion led to a marked reduction in xenobiotic metabolism, with significant suppression of the NRF2 activity in vivo, as identified by IPA upstream regulator analysis and GSEA (**Fig. 6A-B, Sup. Fig. S2I, S2L**). This was accompanied by reduced transcript levels of canonical NRF2 target genes, such as *Nqo1* and *Aldh3a1*, without a change in *Nfe2l2* mRNA expression, the gene encoding NRF2 (**Fig. 6C**).

**Figure 6.**
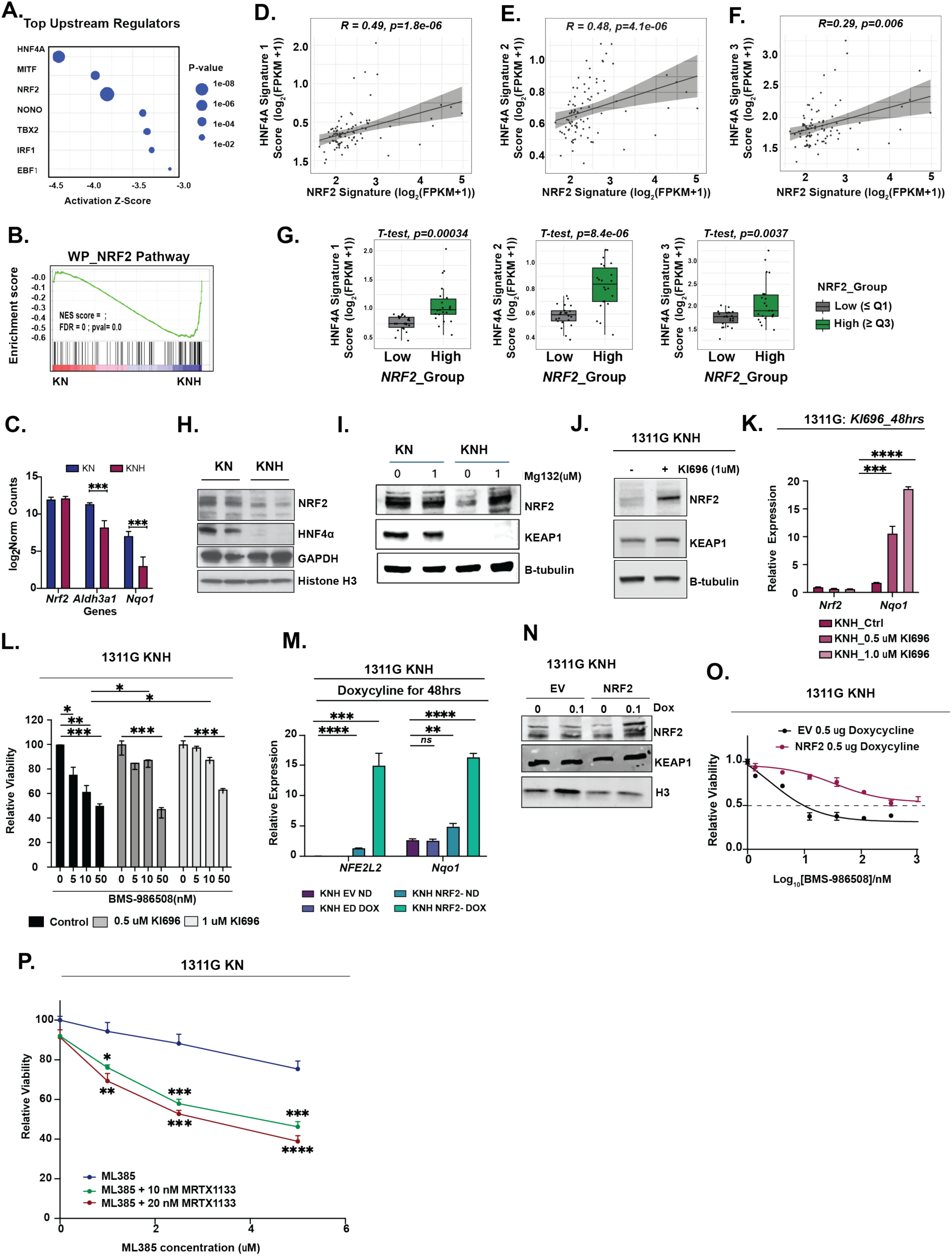
HNF4α impairs response to KRAS inhibition by maintaining NRF2 activity in IMA. **A.** Dot plot showing upstream transcriptional regulators predicted by IPA based on DEGs from in vivo KNH vs KN tumors. **B.** GSEA plot showing enrichment of the Wiki Pathways NRF2 gene signature in DEGs from in vivo KNH vs KN tumors **C.** Log_2_normalized expression of Nrf2 and its downstream targets *Nqo1* (***p = 0.008) and *Aldh3a1* (***p = 0.008) from DESeq2 analysis of bulk RNA-seq comparing KNH to KN in vivo. **D-F.** Spearman correlations between NRF2 activity and HNF4A signature scores across 88 KRAS-mutant NSCLC tumors with wild-type KEAP1 from TCGA. NRF2 activity was quantified using a published NRF2 signatureHNF4A activity was assessed using three independent gene signatures: Signature 1 (**D**), Signature 2 (**E**), and Signature 3 (**F**). Each dot represents a tumor. Spearman correlation coefficient (R) and p-value are shown. Linear regression lines with 95% confidence intervals (gray shaded area) are overlaid. **G.** Boxplots comparing HNF4A signature 1, 2 and 3 activity scores between *NRF2*_Low and *NRF2*_High tumors. All three signatures were significantly elevated in the NRF2_High group. p- values were determined using unpaired two-tailed Student’s *t*-test. **H.** Immunoblot analysis of NRF2 protein levels in KN and KNH tumors FACS sorted from GEMMs 14 weeks post tumor initiation, showing reduced NRF2 expression following *Hnf4a* deletion. **I.** Immunoblot analysis of indicated proteins in 1311G KN and KNH organoids treated with 1 μM MG132 or vehicle for 2 hours **J.** Immunoblot analysis of NRF2 pathway proteins in 1311G KNH organoids following treatment with 1 μM KI696 for 48 hours. **K.** qRT-PCR analysis of *NFE2L2* and *Nqo1* in 1311G KNH organoids treated with increasing doses of KI696 for 48 hours (n = 2; one representative shown). One-way ANOVA: ***p = 0.0003, ****p < 0.0001. **L.** NRF2 activation by KI696 reduces sensitivity of 1311G KNH organoids to BMS-986508. Organoids were pre-treated with KI696 for 7 days, followed by combination treatment with KI696 and BMS-986508 for 3 days. *p < 0.05, **p < 0.01, ***p < 0.001. **M.** qRT-PCR analysis of *NFE2L2* and *Nqo1* in 1311G KNH organoids expressing doxycycline-inducible FLAG-*NFE2L2* or empty vector, treated with 0.1 μg/mL doxycycline for 48 hours. One-way ANOVA: *NFE2L2* (****p < 0.001, ***p = 0.0002); *Nqo1* (**p = 0.0016, ****p < 0.0001). **N.** Immunoblot for NRF2, KEAP1 and H3 in 1311G KNH treated with varying dose of doxycycline for 72 hours. **O.** 1311G KNH Organoids expressing empty vector (EV) or DOX-inducible NRF2 were treated with BMS-986508 ± 0.5 μg/mL DOX for 72 h. IC_50_ increased from 2.5 nM (EV) to 33.7 nM (NRF2). Data represent one of three independent experiments. Error bars represent SEM from technical replicates. **P.** Cell viability in 1311G KN organoids was measured at four ML385 concentrations (0, 1, 2.5, and 5 uM) under three conditions: ML385 alone (blue), ML385 + 10 nM MRTX1133 (green), and ML385 + 20 nM MRTX1133 (red). One-way ANOVA showed significant differences among the groups. p values for Tukey’s multiple comparison test are shown on the plot (*p < 0.05, **p < 0.01, ***p < 0.001, ****p < 0.0001). Each point represents the mean ± SEM of technical replicates and data represent one of three independent experiments.

NRF2 transcriptional activity was recently associated with poor response to KRAS inhibition in the KRYSTAL-1 trial, even in tumors lacking *KEAP1* mutations(Negrao et al., 2025). Because loss of KEAP1 function further amplifies NRF2 signaling, we next turned to TCGA to determine whether HNF4α-NRF2 crosstalk extends to KRAS-mutant, KEAP1-wildtype tumors. We extracted gene expression profiles from 88 TCGA LUAD samples harboring KRAS mutations but lacking KEAP1 alterations, reasoning that these tumors maintain physiologic control of NRF2. Using this dataset, we analyzed gene expression profiles, stratified by an established NRF2 signature(Negrao et al., 2025; Singh et al., 2021) (**Supplemental Table S12).** Using three distinct HNF4α transcriptional signatures derived from human and murine LUAD models, we found that HNF4α activity was significantly enriched in NRF2-high tumors and correlated with NRF2 score (**Fig. 6D-F**). Across all tumors, Signature 1 (R = 0.49, *p* = 1.8e-06), Signature 2 (R = 0.48, *p* = 4.1e-06), and Signature 3 (R = 0.29, *p* = 0.006) showed moderate positive correlations with *NRF2* activity. Consistently, expression of all three signatures was significantly elevated in *NRF2*-High compared to *NRF2*-Low tumors (*p* < 0.001 for all; **Fig. 6G**,). These results confirm that HNF4α transcriptional programs co-occur with high NRF2 activity even in the absence of KEAP1 mutations. In the KRYSTAL-1 cohort, where some tumors carry KEAP1 mutations and thus have constitutively active NRF2, we again observed moderate, significant positive correlations across all three HNF4α signatures and higher signature expression in NRF2 high case (all p < 0.01, **Sup. Fig. S9A-D**). Together, these findings support a conserved, clinically relevant relationship between HNF4α and NRF2 in KRAS mutant LUAD.

To assess whether this transcriptional link extends to protein regulation, we examined the effect of *Hnf4a* deletion on NRF2 protein levels. Immunoblotting revealed reduced NRF2 levels in sorted KNH tumor cells as well as 1311G organoids. NRF2 levels were rescued by proteasome inhibition in vitro, suggesting HNF4α stabilizes NRF2 protein (**Fig. 6H-I**).

We next asked whether the sensitivity of KNH organoids to BMS-986508 is due in part to decreased NRF2 levels. Pharmacologic activation of NRF2 with KI696(Davies et al., 2016) stabilized NRF2 protein and increased *Nqo1* transcript levels, as confirmed by immunoblot and qRT-PCR (**Fig. 6J-K**). KI696 treatment inhibited response to BMS-986508 in 1311G KNH organoids in a dose-dependent manner (**Fig. 6L**). We also engineered 1311G KNH organoids with a doxycycline-inducible *NFE2L2* construct. Upon doxycycline induction, qRT-PCR and western blotting confirmed robust NRF2 expression (**Fig. 6M-N**). Cell viability assays demonstrated that NRF2 re-expression conferred a dose-dependent rescue of BMS-986508 sensitivity relative to empty vector controls, increasing the IC_50_ by >10 fold (**Fig. 6O, Sup. Fig. S9E-F**). Conversely, inhibition of NRF2 with ML385 significantly sensitized 1311G KN organoids to BMS-986508 in a dose-dependent manner, resulting in reduced viability compared to either agent alone (**Fig. 6P**). Together, these findings indicate that diminished NRF2 activity contributes to increased BMS-986508 sensitivity in IMA and suggest this axis as a therapeutic avenue to explore further.

## DISCUSSION

Cancer progression is driven by the remarkable ability of malignant cells to rewire lineage programs in response to environmental and therapeutic pressures(Hanahan, 2022; Marjanovic et al., 2020). In LUAD, invasive mucinous adenocarcinoma (IMA) exemplifies this phenomenon, adopting a gastric epithelial identity following loss of NKX2-1. However, the mechanisms that stabilize this alternative fate and constrain plasticity have remained incompletely defined(Chang et al., 2021; Snyder et al., 2013).

Our study identifies HNF4α as a central enforcer of gastric lineage fidelity in IMA. Positioned within the transcriptional hierarchy downstream of NKX2-1 loss, HNF4α reinforces epithelial differentiation, restricts cellular plasticity, and maintains organized tissue architecture. Mechanistically, HNF4α limits chromatin accessibility to pioneer transcription factors such as FoxA1/2, thereby suppressing the activation of alternative lineage programs associated with neuronal and hepatic fates(Iwafuchi-Doi et al., 2016; Zaret & Carroll, 2011). Upon HNF4α loss, FoxA1/2 gain de novo access to previously inaccessible loci, driving transcriptional reprogramming toward a dedifferentiated state. These findings parallel HNF4α’s known roles in developmental and adult tissues, where it serves as a guardian of epithelial identity(Garrison et al., 2006; Moore et al., 2016) and align with observations in other malignancies where loss of HNF4α correlates with dedifferentiation and lineage switching(Camolotto et al., 2021; Ning et al., 2010; Pelletier et al., 2011). Notably, our findings stand in contrast to our recent work in a distinct hybrid LUAD subtype, where HNF4α was shown to co-regulate both gastric and pulmonary transcriptional programs(Fort et al., 2024). In IMA, however, HNF4α binding is largely restricted to gastric gene loci, reinforcing a more lineage-constrained transcriptional landscape in IMA.

In addition to maintaining bulk epithelial programs, HNF4α preserves the transcriptional identity of a rare tuft-like cell population that integrates gastric features alongside canonical neuronal and immune-associated gene signatures(Haber et al., 2017). While canonical tuft markers remain detectable after *Hnf4a* deletion, expression of noncanonical tuft-associated genes diminishes, suggesting that HNF4α is critical for maintaining the full transcriptional spectrum of tuft cells(Goto et al., 2019; Silverman et al., 2024; Xiong et al., 2022). Given emerging evidence implicating tuft cells in tumor-immune interactions(Goto et al., 2019), further investigation into the functional role of HNF4α-dependent tuft-like programs in IMA progression and immune modulation is warranted.

Although HNF4α reinforces lineage stability in untreated tumors, its role under therapeutic pressure reveals a more complex picture. Intact HNF4α expression prevents cell cycle exit during in vivo KRAS^G12D^ inhibition by maintaining Cyclin D1 levels and pRB phosphorylation (Puyol et al., 2010). In parallel, HNF4α modulates redox balance through stabilization of NRF2, supporting antioxidant programs essential for tumor cell persistence(Motohashi & Yamamoto, 2004; Weiss-Sadan et al., 2023). Loss of HNF4α disrupts both proliferative and redox adaptation mechanisms, resulting in heightened sensitivity to KRAS inhibition(Araujo et al., 2024; Zewdu et al., 2021)

Although our data demonstrate that is an HNF4α targetable vulnerability in IMA, HNF4α loss also dismantles lineage barriers, enabling FoxA1/2-driven activation of alternative programs that may facilitate lineage switching under therapeutic stress(Orstad et al., 2022). Given the established role of FoxA1/2 in driving neuroendocrine transformation in prostate cancer(Wang et al., 2024), it is plausible that IMA tumors lacking HNF4α could escape KRAS blockade through similar mechanisms, raising concerns about histologic evolution and therapeutic resistance. These findings highlight a potential therapeutic paradox: while HNF4α inhibition may enhance initial responses to KRAS inhibition, it could also promote transcriptional evolution and relapse through activation of plasticity programs. This raises the question of whether targets downstream of HNF4α might be more suitable for clinical purposes. We found that NRF2 and HNF4α gene signature scores were strongly correlated in human KRAS-mutant lung tumors from both TCGA, and the KRYSTAL-1 trial, linking this axis to clinically relevant variation in transcriptional state. Similarly, 1311G KN organoids characterized by higher baseline NRF2 displayed greater viability loss upon BMS-986508 treatment once NRF2 was inhibited with ML385. Targeting downstream effectors such as Cyclin D1 or NRF2 may offer a more selective strategy to eliminate DTPs(He et al., 2024). Future studies should assess whether co-targeting cell cycle and redox pathways can enhance the durability of KRAS-directed therapies without inducing dedifferentiation and increased plasticity as potential escape mechanisms.

Together, our findings position HNF4α as a master regulator of lineage fidelity, chromatin accessibility, and therapeutic response in IMA. By integrating developmental, transcriptional, and therapeutic axes, this study provides a conceptual framework for targeting lineage-defining transcription factors in cancer. Strategies that disrupt survival pathways while safeguarding differentiation states may be critical to achieving durable responses in lineage-plastic malignancies such as IMA and other cancer subtypes.

## EXPERIMENTAL MODELS AND SUBJECT DETAILS

### Animal Studies

Mice harboring *Kras^FSF-G12D^* (Young et al., 2011)*, Rosa-FSF-Cre^ERT2^*(Schönhuber et al., 2014), NKX2-1^flox^ (Kusakabe et al., 2006), *Hnf4a*^flox^ (Hayhurst et al., 2001), *R26-CAG-LSL-Sun1-sfGFP-myc (Mo et al., 2015)* and *p53^frt^ (Lee et al., 2012)* have been previously described. All animals were maintained on a mixed 129/B6 background. All experimental mice were between 2 and 5 months of age at intubation. Mice of both sexes were used throughout each study, though the effect of sex on study results was not assessed except for scRNAseq were we used only male mice. Animal studies were approved by the IACUC of the University of Utah, conducted in compliance with the Animal Welfare Act Regulations and other federal statutes relating to animals and experiments involving animals, and adhered to the principles set forth in the Guide for the Care and Use of Laboratory Animals, National Research Council (PHS assurance registration number A-3031-01).

### Husbandry and housing conditions of experimental animals

Experimental mice were either bred in-house or obtained from external sources. Mice were maintained under controlled housing conditions, including regulated temperature, humidity, and light cycles, and were provided with a standardized diet. Genotyping was performed using DNA extracted from ear biopsies to confirm the presence of transgenic or knockout alleles. Mice lacking the relevant alleles were utilized as control littermates or humanely euthanized. Individual mice were identified using ear tags and assigned unique identifiers. To ensure the genetic fidelity of the models, strict handling protocols were implemented, and comprehensive records were maintained throughout the study.

### Primary 3D Organoid cultures

All primary murine (1311G and 429A) and patient derived organoid(HCI_IMA03) organoid cultures were established within Matrigel (Preclinical Research Shared Resource core facility) submerged in recombinant organoid medium for approximately two weeks (Advanced DMEM/F-12 supplemented with 1X B27 (Gibco), 1X N2 (Gibco), 1.25mM nAcetylcysteine (Sigma), 10mM Nicotinamide (Sigma), 10nM Gastrin (Sigma), 100ng/ml EGF (Peprotech), 100ng/ml R-spondin1 (Peprotech), 100ng/ml Noggin (Peprotech), and 100ng/ml FGF10 (Peprotech)). After organoids were established, cultures were switched to 50% L-WRN conditioned media (Miyoshi & Stappenbeck, 2013). Organoid lines were tested periodically for mycoplasma contamination. To maintain organoid cultures mycoplasma free, all culture media were supplemented with 2.5 ug/ml Plasmocin. HEK293T cells were cultured in DMEM/10% FBS (Gibco). All cell lines were tested periodically for mycoplasma contamination. All culture media were supplemented with 2.5 ug/ml Plasmocin to maintain cell cultures mycoplasma free.

## METHOD DETAILS

### Tumor initiation and tamoxifen administration in vivo

Autochthonous lung tumors were initiated by administering viruses via intratracheal intubation. Adenoviral mSPC-FlpO was used to initiate all tumors in this publication. Adenoviruses were obtained from University of Iowa Viral Vector Core.

Tumor-specific activation of Cre^ERT2^ nuclear activity was achieved by intraperitoneal injection of tamoxifen (Sigma) dissolved in corn oil at a dose of 120mg/kg. Mice received 4 intraperitoneal injections over 5 days starting exactly 6 weeks after tumor initiation. Tamoxifen exposure was extended by providing tamoxifen-containing chow (Envigo) for an additional seven days. BrdU incorporation was performed by injecting mice at 40mg/kg (Sigma) intraperitoneally 1 hour prior to tissue collection.

### Histology and immunohistochemistry

All tissues were fixed in 10% formalin overnight and when necessary, lungs were perfused with formalin via the trachea. Organoids were first fixed in 10% formalin overnight and then mounted in HistoGel (Thermo Fisher Scientific). Mounted organoids and tissues were transferred to 70% ethanol, embedded in paraffin, and four-micrometer sections were cut. Immunohistochemistry (IHC) was performed manually on Sequenza slide staining racks (Thermo Fisher Scientific). Sections were treated with Bloxall (Vector Labs) followed by Horse serum (Vector Labs) or Rodent Block M (Biocare Medical), primary antibody, and HRP-polymer-conjugated secondary antibody (anti-Rabbit, Goat and Rat from Vector Labs; anti-Mouse from Biocare). The slides were developed with Impact DAB (Vector Labs) and counterstained with hematoxylin. Slides were stained with antibodies to NKX2-1 (1:2000, Abcam, EP1584Y), CC3 (CST 9664S 1:400), GFP (1:200, CST, 2956S), FoxA1 (1:4000, Abcam, 10881-14), FoxA2 (1:1200, Abcam, 4466), HNF4a (1:500, CST, C11F12), pro-SPC (1:4000, Abcam, EPR19839), Galectin 4 (1:200, R and D Systems, AF2128), Muc5AC (1:100, Abnova, SPM488), p63 (1:500, CST, D2K8X), Cytokeratin 5 (1:200, Abcam, EP1601Y), Gastrokine 1 (1:50, Abnova, 2E5), pERK1/2 (1:500, CST, D13.14.4E), TFF1 (1:100, Origene, TA382458), Pepsinogen C (1:100, Sigma, HPA031718), MCM2 (1:1000, CST 4007), pRBS807(1:1000; CST D20B12), NRF2(1:100, CST 12721). Images were taken on a Nikon Eclipse Ni-U microscope with a DS-Ri2 camera and NIS-Elements software. Using NIS-Elements software, tumor Burden, BrdU quantitation and histological analyses were performed on hematoxylin and eosin-stained and IHC-stained slides. All histopathologic analysis was performed by a board-certified anatomic pathologist (E.L.S.).

### Ethics for Patient derived Xenografts

Tumor acquisition and experimental usage was approved by the University of Utah IRB (89989 and 10924) and the animal studies were approved by the University of Utah Institutional Animal Care and Use Committee (17-06008).

### Establishing Primary Murine and Human LUAD organoids

**429A** - 10 weeks after tumor initiation in KNH (*Kras^FSF-G12D/+^*^;^ *Rosa26^FSF-CreERT2^; Nkx2-1^F/F^; Hnf4a^F/F^)* mice, tumor bearing mice were euthanized and lungs were isolated. Whole lung full of microscopic tumors were minced under sterile conditions and digested at 37°C for 30 min with continuous agitation in a solution of Advanced DMEM/F12 containing the following enzymes: Collagenase Type I (Thermo Fisher Scientific, 450U/ml), Dispase (Corning, 5U/ml), DNase I (Sigma, 0.25mg/ml). Enzymatic reactions were stopped by addition of cold Advanced DMEM/F-12 with 10% FBS. The digested tissue was repeatedly passed through a 20-gauge syringe needle, sequentially dispersed through 100 mm, 70 mm, and 40 mm cell strainers, and treated with erythrocyte lysis buffer (eBioscience) to obtain a single cell suspension.

Organoid cultures were established by seeding 1×10^5^ tumor cells in 50ul of Matrigel (Corning) and plated in 24-well plates. Matrigel droplets were overlaid with recombinant organoid medium as described above (Cell lines and primary cultures). Cultures were switched to 50% L-WRN conditioned media two weeks after organoid establishment. Organoid cultures were screened via immunohistochemistry and qPCR, and lines that expressed both HNF4α and NKX2-1 were selected for subsequent analysis (Pleguezuelos-Manzano et al., 2020) to 429A organoid lines.

**1311G#5**- We established the tumor from KNH (*Kras^LSL-G12D/+^; Nkx2-1^F/F^; Hnf4a^F/F^)* using a lox-stop Cre viral delivery system. However, due to the poor recombination efficiency of viral Cre, we selectively harvested tumors that were *Nkx2-1* negative but *Hnf4aα* positive, as described above. By delivering Cre virus in vitro, we generated isogenic organoid lines that were either HNF4α-positive (KN) or HNF4α-negative (KNH).

The full names of the 3D cell lines mentioned in this paper are as follows: **1311G** refers to “Mouse 1311, Tumor G, Subclone 5,” and **429A** denotes “Mouse HED 429 Tumor, Subclone A.” These names have been abbreviated for clarity and brevity.

**HCI_IMA03** - Patient-derived organoids were established from a surgically resected primary invasive mucinous adenocarcinoma (IMA) of the lung obtained from a patient at Huntsman Cancer Institute under institutional review board (IRB)-approved protocols. Tumor tissue was minced under sterile conditions and digested at 37°C for 30 minutes with continuous agitation in Advanced DMEM/F12 containing Collagenase Type I (Thermo Fisher Scientific, 450 U/ml), Dispase (Corning, 5 U/ml), and DNase I (Sigma, 0.25 mg/ml). Enzymatic reactions were stopped with cold Advanced DMEM/F12 supplemented with 10% FBS. Tissue was further dissociated by repeated passage through a 20-gauge syringe needle, filtered through 100 µm, 70 µm, and 40 µm cell strainers, and erythrocyte lysis buffer (eBioscience) was applied to remove red blood cells. The resulting single-cell suspension was embedded in 50 µl Matrigel (Corning) and seeded into 24-well plates. After Matrigel polymerization, wells were overlaid with recombinant organoid medium as described above (Cell lines and primary cultures). Cultures were maintained in 50% L-WRN conditioned media following initial outgrowth. Organoids were screened by histology and qPCR for expression of HNF4α and NKX2-1, and positive clones were expanded for downstream analysis.

### In vitro 4-hydroxytamoxifen (4OHT) treatment

429A organoid was transiently treated with 2mM 4OHT (Cayman Chemical Company, dissolved in 100% Ethanol) or vehicle for 48hr to activate Cre^ERT2^ nuclear activity and generate isogenic pairs that either positive or negative for HNF4α.

### Generating a single cell suspension from organoid cultures

Matrigel droplets containing organoids were broken down via repeated pipetting in Cell Recovery Solution (Corning, 500 µl per Matrigel dome). Cell Recovery Solution containing organoids was transferred to sterile conical tubes and submerged in ice for 20-30 min before centrifugation at 4°C (300-500G). Cell Recovery Solution supernatant was removed, and the cell pellet was washed in 1X PBS. Cells were then resuspended in pre-warmed TrypLE Express Enzyme (Thermo Fisher Scientific) and incubated for 10 min at 37°C. TrypLE reaction was quenched via dilution with cold Splitting Media (Advanced DMEM/F-12 [Gibco], 10 mM HEPES [Invitrogen], 1X Penicillin-Streptomycin-Glutamine [Invitrogen]). Cells were centrifuged and then resuspended in a pre-warmed DNase solution (L-WRN media supplemented to a final concentration of 200U/ml DNase [Worthington], 2.5 mM MgCl2, 500 mM CaCl2) and incubated for 10 min at 37°C. Cells were centrifuged and washed in PBS before use.

### Immunoblotting

Protein was extracted by lysing cells on ice for 20 mins in RIPA buffer (50 mM Tris HCl pH 7.4, 150 mM NaCl, 0.1% (w/v) sodium dodecyl sulfate, 0.5% (w/v) sodium deoxycholate, 1% (v/v) Triton X-100) plus Pierce Protease Inhibitor (PPI, ThermoFisher Scientific, A32959). Cellular debris was pelleted for 15 min at 4°C and protein concentration was quantitated with the Pierce Bradford Protein Assay Kit (ThermoFisher Scientific, 23200). A total of 20μg of protein lysates were separated on Tris-Glycine precast gels (SMOBIO, QP4510) and transferred to nitrocellulose membrane (ThermoFisher Scientific, 88018). Membranes were probed overnight with antibodies against Tubb(1:1000, DSHB-E7), FoxA1 (1:1000, Abcam 23738), FoxA2 (1:1000, Abcam 108422), NKX2-1 (1:2000, Abcam 133638), HNF4α (1:1000, CST C11F12 or 1:1000, PP-H1415-0C), pERK (1:1000, CST 4370), total ERK (1:2000, CST 4695), KEAP1(1:1000, CST 8047), GAPDH(1:1000, CST 5174), NRF2(1:1000, CST D1Z9C) and Vinculin (1:20000, Abcam 129002). After incubation with primary antibodies, membranes were incubated with either IRDye 800CW Goat anti-Rabbit IgG or IRDye 680RD Goat anti-Mouse IgG secondary antibodies (1:15,000 dilution; LI-COR Biosciences) and imaged using a LI-COR Odyssey CLx scanner and Image Studio software, or alternatively with HRP-conjugated secondary antibodies (1:5,000; Cell Signaling Technology) and detected by enhanced chemiluminescence (ECL) using SuperSignal West Femto Maximum Sensitivity Substrate (Thermo Fisher Scientific) and imaged with a Bio-Rad ChemiDoc MP system. Densitometric quantification was performed using Image Studio or Image Lab software as appropriate.

### Subcutaneous cell line xenografts

KN and KNH cells were collected and mixed with Matrigel 1:1 by volume for subcutaneous xenograft experiments. Cells were subcutaneously injected into the flanks of NOD/SCID-gamma chain deficient mice (NSG) (∼0.4 million cells/flank for 1311G and 1 million cells/flank for 429A). Tumors were allowed to establish, and their dimensions were measured using calipers. Tumor volume was calculated using the formula (L x W^2^)/2 where (L) is the tumor’s length and (W) its width. Tumor volumes were monitored weekly, with measurements taken every other day. Mice were euthanized when any tumor within the cohort exceeded a volume of 1000 mm³.

### Lentiviral production and transduction

Lentivirus were produced by transfection of HEK-293T cells with TransIT-293 (Mirus Bio). Packaging vectors Δ8.9 (gag/pol) and VSV-G were used for lentiviral production. Supernatant was collected at 48- and 72-hours post-transfection, centrifuged, and filtered using 0.45um filter units before storing long term at −80°C.

To achieve stable transduction of organoids, cultures were dissociated into single-cell suspensions as previously described (see “Generating a Single-Cell Suspension from Organoid Cultures”). The cells were resuspended in a 1:1 mixture of 50% L-WRN and thawed lentivirus-containing supernatant. Polybrene was added to a final concentration of 8 µg/mL, and the cells were incubated for 24 hours. Following incubation, the cells were pelleted, re-embedded in Matrigel, and seeded. After 72 hours, antibiotic selection (Puromycin = 5µg/mL or Blasticidin = 10µg/mL) was done for at least one week to establish stable lines for downstream assays.

### BMS-986508 treatment in autochthonous and allograft mouse models

BMS-986508 was provided by BMS and formulated at 10% (w/v) Captisol in 50 mM citrate buffer, pH 5.0, by mixing Captisol and citrate buffer, adjusting pH with NaOH, and dissolving drug by vortexing and heating to 37°C prior to use. For allograft studies, NSG mice were subcutaneously implanted with either KN or KNH 1311G tumor organoids suspended in Matrigel (Corning). Tumor-bearing mice were randomized once tumors reached ∼150 mm³. BMS-986508 or vehicle control (10% Captisol in 50 mM citrate buffer) was administered by daily intraperitoneal (i.p.) injection at a dose of 10 mg/kg for 7 days, followed by dose escalation to 30 mg/kg for an additional 14 days. Mice were then weaned off drug for a 14-day observation period prior to sacrifice.

For autochthonous tumor models, KN and KNH mice were administered tamoxifen to initiate tumor development as described above. BMS-986508 was delivered by daily i.p. injection at 30 mg/kg in 10% Captisol in 50 mM citrate buffer (pH 5.0), beginning at 12 weeks post-tumor initiation, for a total duration of 14 consecutive days prior to sacrifice. All animal experiments were conducted under protocols approved by the University of Utah Institutional Animal Care and Use Committee (IACUC).

### PrestoBlue Cell Viability Assays

Organoids were broken down into a single cell suspension as described above (*Generating a single cell suspension from organoid cultures*) and seeded at equal density with 5ul of Matrigel per well in a solid wall, transparent bottom 96 well plate with 100ul of LWRN per well whilst chronically For growth and drug treatment assays, 429A cells were pretreated with 4OHT (or ethanol as a control) for 48 hours before seeding, while chronically deleted *Hnf4a* 1311G cells were used when necessary. Baseline measurements were performed one day post-seeding by adding 10 µL of PrestoBlue HS Cell Viability Reagent (Invitrogen) to each well, followed by incubation at 37°C for 30 minutes. Fluorescent emission was quantified using a Synergy HTX plate reader (Excitation: 528/20 nm, Emission: 590/20 nm). After measurement, the PrestoBlue reagent was removed, wells were washed with warm PBS, and fresh LWRN media was added. Measurements were then taken every other day for growth assays or after 72 hours for drug treatment experiments.

### RNA extraction, cDNA synthesis, and qPCR

RNA was isolated via Trizol-chloroform extraction followed by column-based purification. The aqueous phase was brought to a final concentration of 35% ethanol, and RNA was purified using the PureLink RNA Mini Kit (ThermoFisher Scientific) according to the manufacturer’s specifications.

cDNA was synthesized from Trizol-extracted RNA using LunaScript RT SuperMix (NEB, M3010) according to manufacturer’s specifications. qPCR was performed on cDNA using Luna Universal Probe qPCR Master Mix (NEB, M3004) according to manufacturer’s specifications, and 35 cycles were used for the denaturation and extension steps. Transcript levels were normalized to PPIA and quantitated by the ΔΔCt method. All Taqman probes utilized are listed in the key resources table.

### RNA sequencing

#### In vivo RNA sequencing

14 weeks after tumor initiation, KN and KNH mice were euthanized, and the ribcage was dissected to reveal the trachea and heart. Cardiac perfusion of the pulmonary vasculature was performed using PBS until the lungs turned pale. Lungs were then removed, digested, and filtered as previously described (Establishing primary murine LUAD organoids) to obtain a single cell suspension. Samples were resuspended in FACS buffer with DAPI. Cells were sorted on the BD FACSAria with the 85 mm nozzle to obtain a GFP-positive, DAPI-negative population. Samples were sorted into 1 ml of cold PBS with 10% serum. After sorting, cells were centrifuged for 10 min at 4 C (300G) and resuspended in 1 ml of Trizol. RNA was isolated via Trizol-chloroform extraction and the PureLink RNA Mini kit as previously described (In vitro RNA-seq). Library preparation was performed using the NEBNext Ultra II Directional RNA Library Prep with rRNA Depletion Kit for mouse. Sequencing was performed using the Illumina NovaSeq 6000 (150 x 150 bp paired-end sequencing, 25 million reads per sample).

#### In vitro RNA sequencing

RNA was collected from 2 biological replicates of the following conditions: 429A KN/KNH isogenic organoid cultures (2 weeks following 4OHT treatment), 1311G KN/KNH isogenic organoids (>4 weeks following Ad5CMV -Cre treatment) and 1311G KN/KNH BMS-986508 treated cells (48h of treatment with vehicle or 4nM BMS-986508). Cells were collected directly into Trizol and stored at −80°C until purification. For organoids, 4 confluent 20μL Matrigel domes were collected per sample.

RNA was isolated via Trizol-chloroform extraction followed by column-based purification. The aqueous phase was brought to a final concentration of 35% ethanol, and RNA was purified using the PureLink RNA Mini kit according to the manufacturer’s instructions (ThermoFisher Scientific). Library preparation was performed using the NEBNext Ultra II Directional RNA Library Prep with poly(A) mRNA isolation. Sequencing was performed using the Illumina NovaSeq 6000 (150 x 150 bp paired-end sequencing, 25 million reads per sample).

#### RNA-seq data processing and analysis

The mouse mm10 (or mm39 for 3154 BMS-986508+/-Hnf4a) and gene feature files were downloaded from Ensembl and a reference database was created using STAR version 2.7.6a (Dobin et al., 2012). Optical duplicates were removed from NovaSeq runs via Clumpify v38.34 (Bushnell, 2014). Reads were trimmed of adapters and aligned to the reference database using STAR in two-pass mode to output a BAM file sorted by coordinates. Mapped reads were assigned to annotated genes using feature Counts version 1.6.3. (Liao et al., 2019). Raw counts were filtered to remove features with zero counts and features with five or fewer reads in every sample. DEGs were identified using the hciR package (https://github.com/HuntsmanCancerInstitute/hciR) with a 5% false discovery rate and DESeq2 version 1.34.0 (Love et al., 2014). GSEA-Preranked was run with the differential gene list generated from DESeq2 and the following gene sets: c2, c5, c8 and Hallmarks gene sets from MSigDB(Liberzon et al., 2011; Subramanian et al., 2005). Gene sets smaller than 15 and larger than 500 were excluded from analysis.

### Single-Cell RNA Sequencing

#### Sample Preparation

KN(n=2) and KNH (n=2) mice were administered tamoxifen 6 weeks post-intubation for a duration of 2 weeks (one week of IP tamoxifen IP dose followed by one week on tamoxifen chow). 6 weeks later, single-cell suspensions were prepared as follows: Lungs and heart were perfused with PBS, and whole lung filled with microscopic tumors were dissected and dissociated into single cells as outlined in the “Establishing Primary Murine LUAD Cell Lines and Organoids” protocol. Samples were resuspended in PBS +1% BSA buffer with DAPI. Cells were sorted on the BD FACSAria cell sorter with the 85 mm nozzle to obtain a GFP-positive, DAPI-positive population. After sorting, cells were centrifuged for 10 min at 4 C (300G) and resuspended in PBS with 1% BSA for library preparation on the same day.

#### Library Preparation and Sequencing

The scRNA-seq libraries were generated using the 10x Genomics Chromium Single Cell Gene Expression Solution with 3’ chemistry (version 3, PN-1000075). This process was conducted at the High-Throughput Genomics Shared Resource, Huntsman Cancer Institute, University of Utah. Single-cell suspensions, filtered through a 40 µm strainer, were assessed for viability and cell count using Countess II (Thermo Scientific) and adjusted to a target recovery of 10,000 cells. The suspension was loaded into Chromium Single Cell A (PN-120236), where Gel Beads-in-Emulsion (GEMs) were formed. Reverse transcription synthesized cDNA from barcoded mRNA within GEMs, followed by subsequent library preparation steps, including A-tailing, end repair, adaptor ligation, and indexing. Library quality was assessed with Agilent D1000 ScreenTape and quantified via qPCR using KAPA Biosystems Library Quantification Kit for Illumina Platforms (KK4842). Libraries were normalized and sequenced on a NovaSeq 6000 in paired-end mode (2

× 150).

### scRNA-seq Data Processing and Analysis

#### Demultiplexing and Alignment

Demultiplexing of scRNA-seq data from KN (n=2) and KNH (n=2) tumors was performed using Cell Ranger (mkfastq version 3.1.0) to generate fastq files. Reads were aligned to the mouse genome (mm10), supplemented with references for CRE-ERT2 and HNF4α exons, using Cell Ranger count (version 3.1.0). The expected cell count was set to 10,000 per library. For KN samples, approximately 6,497 cells were captured, with an average of 36,677 reads per cell and a median of 3,329 genes per cell. For KNH samples, approximately 5,817 cells were captured, with an average of 41,861 reads per cell and a median of 3,641 genes per cell. Additional details of the primary Cell Ranger data processing can be found at: <u>link to 10x</u>

#### Quality Control and Clustering

Single cell expression data was subjected to common Seurat workflows for initial quality control and clustering (https://satijalab.org/seurat/articles/pbmc3k_tutorial.html). Specifically, Seurat (version 4) workflows were employed for QC and clustering. Cells with feature counts <500 or >7,500 and mitochondrial content >10% were excluded. Data were normalized, scaled, and dimensionally reduced using PCA (25 PCs with a resolution of 1). A KNN graph was constructed, and clusters were identified using the Louvain algorithm. UMAP embedding revealed 17 clusters which included some immune cells that were captured. High-quality tumor cells from KN and KNH were identified, sub-setted, and reclustered based on 19 PCs. Tumor cell barcodes used for downstream analyses are provided in Supplemental Table S10. To identify KNH complete recombinants, expression of the floxed fourth and fifth exons of *Hnf4a* were quantified.

#### Differential Gene Expression and Scoring

Differentially expressed genes in UMAP clusters were identified using Seurat’s FindMarkers function. Gene signature scores for published gene signatures including various additional cell types and states (MsigDB) used in this paper were calculated using the AddModuleScore function. Published gene lists for these scores are available in referenced papers. CytoTRACE(Gulati et al., 2020) scores, ranging from 0 (most differentiated) to 1 (least differentiated), were computed using the CytoTRACE2 R package (v1.0.0).

#### Data Imputation

The ALRA method was applied to impute low-detection genes in the scRNA-seq dataset, preserving biological zeros and utilizing low-rank approximation(Linderman et al., 2022).

### Chromatin immunoprecipitation sequencing

#### In vitro organoid ChIP-seq

For all organoid ChIP-Seq experiments, two 24-well plates (approximately 4 - 8 million cells) were collected in Cell Recovery Solution (500 ml per dome). For *Hnf4a* deletion studies, 429A organoids were treated with 4-hydroxytamoxifen (4OHT) or ethanol as a control 48 hours to induce *Hnf4a* deletion. Additionally, we utilized 1311G organoids that had been previously treated with Ad5CMV-Cre (to delete *Hnf4a*) invitro. Organoids were submerged in ice for 30 min before centrifugation at 4 C (300G). Cells were washed three times in cold PBS. On the second wash, PBS was supplemented with DNase solution (containing a final concentration of 200U/ml DNase [Worthington], 2.5 mM MgCl2, 500 mM CaCl2). After the third wash, organoids were resuspended in 5 ml of 2 mM DSG buffer (1X PBS, 1 mM MgCl2) and rotated at room temperature for 35 min. Formaldehyde was then added to a final concentration of 1% and cells were crosslinked for 10 min. The cross-linking reaction was stopped with the addition of glycine to a final concentration of 125mM. Cells were washed with cold PBS, then frozen at −80. Cell pellets were thawed on ice for 5 minutes then lysed in 1 mL of Farnham lysis buffer (5 mM PIPES pH 8.0, 85 mM KCl, 0.5% NP40) with Pierce Protease Inhibitor (PPI, ThermoFisher Scientific, A32959). Samples were centrifuged at 4°C (1000G) then resuspended in 1mL of RIPA lysis buffer (1X PBS, 1% NP40, 0.5% sodium deoxycholate, 0.1% sodium dodecyl sulfate) with PPI. Chromatin was sonicated with a QSonica Q800R (pulse: 30 son/30 s off; sonication time: 20 minutes; amplitude: 70%). After sonication, samples were centrifuged at 17,000 x g for 15 minutes and an input was collected from each sample of sheared chromatin. Chromatin from each sample was then immunoprecipitated overnight with 5ug of the following antibodies (per sample) premixed with Protein G Dynabeads (for mouse antibodies; ThermoFisher Scientific, 10004D) or Protein A Dynabeads (for rabbit antibodies; ThermoFisher Scientific, 10002D): HNF4α (Perseus Proteomics PPH1415), FoxA1 (Abcam, ab170933) and FoxA2 (CST 8186). The next day, bound chromatin was washed 5 times with LiCl Wash Buffer (100 mM Tris pH 7.5, 500 mM LiCl, 1% NP-40, 1% sodium deoxycholate) and crosslinks were reversed by incubating samples in IP Elution Buffer (1% SDS, 0.1% NaHCO_3_) overnight at 65°C. ChIP DNA was purified using Zymo ChIP DNA Clean and Concentrator Kit (Zymo, D5205).

Library preparation was performed using the ChIP-Seq with NEBNext DNA Ultra II library prep kit using Unique Molecular Indexes (UMIs). Sequencing was performed using the Illumina NovaSeq 6000 (150 x 150 bp paired-end sequencing, 25 million reads per sample).

#### In vivo nuclei ChIP-seq

14 weeks after tumor initiation, KN and KNH tumor-bearing mice were euthanized, and the ribcage was dissected to reveal the trachea and heart. Lungs were perfused with cold PBS, removed, and snap frozen in liquid nitrogen. Flash-frozen lungs were minced on ice in 2 ml of ice-cold PBS for 3 - 5 min. Minced lungs were dounced 10 times with a large 7 ml homogenizer (Wheaton, item #23ND78) and centrifuged for 5 min at 4 C (300G). Tissue was crosslinked in 2 mM DSG buffer and 1% formaldehyde as previously described (In vitro organoid ChIP-seq). After fixation, lung samples were resuspended in 2 - 3 ml of TST buffer with protease inhibitors for 5 min. During this time, samples were dounced an additional 5 times in a small 2 ml homogenizer (Wheaton, item #23ND70) to extract nuclei. Lysis was quenched with 5 ml of 1X ST buffer plus protease inhibitors. Nuclei were washed in cold PBS and resuspended in 5 - 10 ml cold PBS supplemented with DAPI and protease inhibitors. Nuclei suspension was then sequentially filtered through a 70 mm and 35 mm cell strainer. Before sorting, nuclei were evaluated on a fluorescence microscope to assess for GFP positivity and nuclear integrity. Nuclei were sorted using BD FACSAria with the 85 mm nozzle for GFP-positive, DAPI-positive nuclei. Samples were sorted into 1 ml of cold PBS with 1% BSA and 10X protease inhibitor. Approximately 10 million nuclei were sorted for ChIP-Seq for TF ChIP-Seq experiments. After sorting, nuclei were pelleted for 10 min at 4 C (500G). Samples were then resuspended in Chromatrap hypotonic and lysis buffers as previously described (In vitro organoid ChIP-seq) and sonicated with QSonica Q800R (pulse: 30s on /30s off; sonication time: 20 min; amplitude: 70%). Chromatin was immunoprecipitated with antibodies premixed with Protein A Dynabeads: HNF4α (Perseus Proteomics PPH1415), FoxA1 (Abcam, ab170933, rabbit monoclonal, 5 mg/ChIP) and FoxA2 (CST, D56D6, rabbit monoclonal, 5 mg/ChIP). ChIP-Seq libraries were prepped and sequenced as previously described (In vitro organoid ChIP-seq).

#### ChIP-Seq data processing and analysis

Fastq alignments were pre-processed with the merge_umi_fastq application from the UMIScripts package (https://github.com/HuntsmanCancerInstitute/UMIScripts) to associate the UMI sequence, provided as a third Fastq file, into the read comment. Reads were aligned using Bowtie2 v2.2.9(Langmead & Salzberg, 2012) to the standard chromosomes of the mouse genome (version mm10) or the human genome (version hg38). Duplicate alignments based on the UMI code were removed using the bam_umi_dedup application (UMI Scripts) allowing for 1 mismatch. Peaks were called using MACS2 v2.2.7 (Zhang et al., 2008) with a significance of q-value < 0.01. Coverage tracks were generated with MACS2 as Reads Per Million. Input libraries were obtained from all cell line and organoid samples and were used as controls for each ChIP-Seq experiment. All ChIP-Seq experiments were performed in biological duplicates. Peaks called in both biological replicates were identified using Bed tools v2.28.0 (Quinlan & Hall, 2010) with a 1-bp minimum overlap to generate a consensus list of peaks for downstream analysis. Genomic annotation of binding sites was performed using HOMER (Heinz et al., 2010). Motif analysis was performed on 100-bp regions surrounding the summit of identified peaks using the HOMER package.

Differential ChIP-Seq peaks were identified using the Diffbind package v3.4.11 (https://bioconductor.org/packages/release/bioc/html/DiffBind.html) with a q-value cutoff < 0.05 using DESeq (Love et al., 2014) for the analysis method. Motif finding for differential ChIP-Seq peaks was performed using HOMER using the entire 400bp differential peak region output by DiffBind. Heat maps and profile plots were generated using deeptools v3.5.1(Ramírez et al., 2016). Pathway analysis was performed on annotated differential ChIP-Seq peaks using Enrichr (Edward Y. Chen et al., 2013).

#### Integration of HNF4α ChIP-seq and H3K27ac HiChIP Data

Previously published H3K27ac HiChIP data from KN GEMM tumors (Gillis et al., 2023) were integrated with HNF4α ChIP-seq dataset from KN GEMM tumors to identify enhancer- and promoter-associated chromatin interactions. High-confidence loops (FDR < 0.01, HiChIPcounts > 2) were filtered to remove duplicates and adjacent anchors corresponding to the same transcriptional unit.

For each retained loop, anchor regions were compared to HNF4α peak coordinates (from MACS2 peak calls) to identify direct overlaps. The gene-proximal anchor (within the transcribed region or promoter) was inferred using loop-associated gene annotations. The distal anchor was designated as the putative enhancer. Anchors overlapping HNF4α peaks were flagged as HNF4A-bound, and each loop was classified as promoter, enhancer, or both, depending on the location of HNF4α binding. Genes with both promoter- and enhancer-bound loops were annotated as both. Final annotations included loop coordinates, HNF4α binding status, loop strength (HiChIPcounts), FDR, and associated gene identity.

#### NRF2 and HNF4A Signature Scoring and Stratification Analysis

Gene expression data from 88 NSCLC lung adenocarcinoma (LUAD) tumors with confirmed *KRAS* mutations and wild-type *KEAP1* status were obtained from The Cancer Genome Atlas (TCGA). Normalized transcript abundance (log₂(FPKM + 1)) was used after removing genes with duplicated names or zero expression across all samples. HNF4A and NRF2 signature scores were computed by averaging the log₂-transformed expression of genes in each signature, retaining only those present in the dataset. Signature 1 comprised the top 100 DEGs upregulated in KN GEMM tumors; Signature 2 included 100 DEGs induced upon HNF4A overexpression in H2122 cells; Signature 3 was derived from hybrid human–mouse LUAD models(Fort et al., 2024). NRF2 activity was assessed using a published gene set utilized in the KRYSTAL study(Negrao et al., 2025; Singh et al., 2021). Samples were stratified into NRF2_Low (≤Q1) and NRF2_High (≥Q3) groups based on NRF2 scores, excluding intermediate cases. Two-tailed unpaired Student’s t-tests (α = 0.05) were used to compare HNF4A scores between NRF2_Low and NRF2_High tumors. Spearman correlations between NRF2 and HNF4A activity scores were computed across all 88 tumors.

For comparison, gene expression data from 68 KRAS-mutant NSCLC patients enrolled in the KRYSTAL-1 clinical trial (NCT03785249)(Negrao et al., 2025) were obtained from the supplementary materials of a previously published study(Tong et al., 2024). Genes with zero counts across all samples and those with CPM < 1 in >50% of samples were excluded. Remaining counts were normalized using edgeR (v3.36.0), and log₂CPM values were computed with a prior count of one. HNF4A and NRF2 signature scores were calculated as above. Tumors were stratified into NRF2_Low and NRF2_High based on quartiles, and two-tailed unpaired t-tests were used to compare HNF4A activity scores between groups. Spearman correlation analyses were also performed across the full cohort. Boxplots and scatter plots were generated in R using ggplot2 (via ggpubr v0.4.0 and cowplot v1.1.1), with a single least-squares regression line, 95% confidence intervals, and annotated Spearman R and p values.

## QUANTIFICATION AND STATISTICAL ANALYSIS

All graphing and statistical analysis was performed with PRISM software or R, with all graphs showing mean and standard deviation or standard error. The statistical details can be found in the corresponding figure legend. All NGS statistical analysis was performed according to published pipeline protocols cited, with a statistical significance cutoff of padj<0.05.

## Supporting information

HED Supp Figures

## ACKNOWLEDGEMENTS

We are grateful to members of the Snyder lab for their helpful suggestions and feedback. We thank B. Dalley for sequencing expertise, J. Marvin for FACS support and O. Allen for expertise in scRNA-seq. E.L.S. was supported by grants from the NIH (R01CA212415, R01CA240317, and R01CA237404), the American Lung Association (LCD-821670), and institutional funds from the Department of Pathology and Huntsman Cancer Institute/Huntsman Cancer Foundation, University of Utah. M.G was supported by the Lung Cancer Research Foundation. Y.M. was supported by R01CA240317. B.T.S. was supported by R01CA289704. The graphical abstract for this publication was created with Biorender.com. Research reported in this publication utilized shared resources at the University of Utah, including the High Throughput Genomics, Bioinformatics, Flow Cytometry, Biorepository and Molecular Pathology cores, and The Center for High Performance Computing. Core facility support was provided by the National Cancer Institute of the National Institutes of Health under award number P30CA042014. Work in the Flow Cytometry Core Facility was additionally supported by the National Center for Research Resources of the NIH under award number 1S20RR026802-1. The content is solely the responsibility of the authors and does not necessarily represent the official views of the NIH.

## AUTHOR CONTRIBUTIONS

H.E.D and E.L.S. designed experiments. H.E.D., Y.S.G. and M.G. performed experiments. H.E.D. and E.L.S. analyzed data. E.L.S. performed histopathologic review. H.E.D. and E.L.S. wrote the manuscript with input and analysis from B.T.S. Y.M. provided data. All authors discussed results, reviewed and revised the manuscript.

## SUPPLEMENTAL INFORMATION

**Table S1.** ChIP-seq peaks and gene annotations for GEMM (KN and KNH), 1311G, 429A and HCI_IMA03, related to Figures 2, S2 and S6

**Table S2.** Raw counts, normalized counts and differential expressed genes (DEGs) in GEMM (KN and KNH) tumors, related to Figures 2 and S3.

**Table S3.** Integration of HNF4α ChIP-seq with H3K27ac HiChIP in KN GEMM tumors, related to Figure 2.

**Table S4.** GSEA cell-type (C8) analysis of DEGs in GEMM (KN and KNH) tumors, related to Figures 2 and S3.

**Table S5.** Raw counts, normalized counts and DEGs in 429A murine organoids, related to Figure S3.

**Table S6.** Raw counts, normalized counts and DEGs in 1311G murine organoids, related to Figure S3.

**Table S7.** GSEA cell-type (C8) analysis of DEGs in 429A and 1311G murine organoids, related to Figure S3.

**Table S8.** Halmark, KEGG and Reactome analysis of DEGs from GEMM (KN and KNH) tumors, related to Figure S3.

**Table S9.** GSEA cell-type (C8) analysis of pooled DEGs from 429A and 1311G murine organoids, related to Figure S3.

**Table S10.** scRNA-seq tumor cell barcodes, cluster associations, cluster-specific DEGs, HNF4α exon counts, and pathway analysis between Group A and B, related to Figures 3, S4 and S5.

**Table S11.** Differentially bound FoxA1/2 peaks and gene annotations in GEMM (KN and KNH) tumors and 1311G organoids, related to Figures 4 and S7.

**Table S12.** Published NRF2 gene signature, HNF4A Signature 1-3 and TCGA gene expression data for KRAS mutant, KEAP1 wild type tumors, related to Figures 6 and S9.

## RESOURCE AVAILABILITY

### Lead contact

Further information and requests for resources and reagents should be directed to and will be fulfilled by the lead contact, Eric Snyder

### Materials availability

Murine organoid lines and Patient derived organoid generated in this study are available upon request.

