## Supplementary material for "HNF4α controls growth, identity and response to KRAS inhibition of invasive mucinous adenocarcinoma of the lung": HED Supp Figures

### Supplement 1, related to main Figure 1.

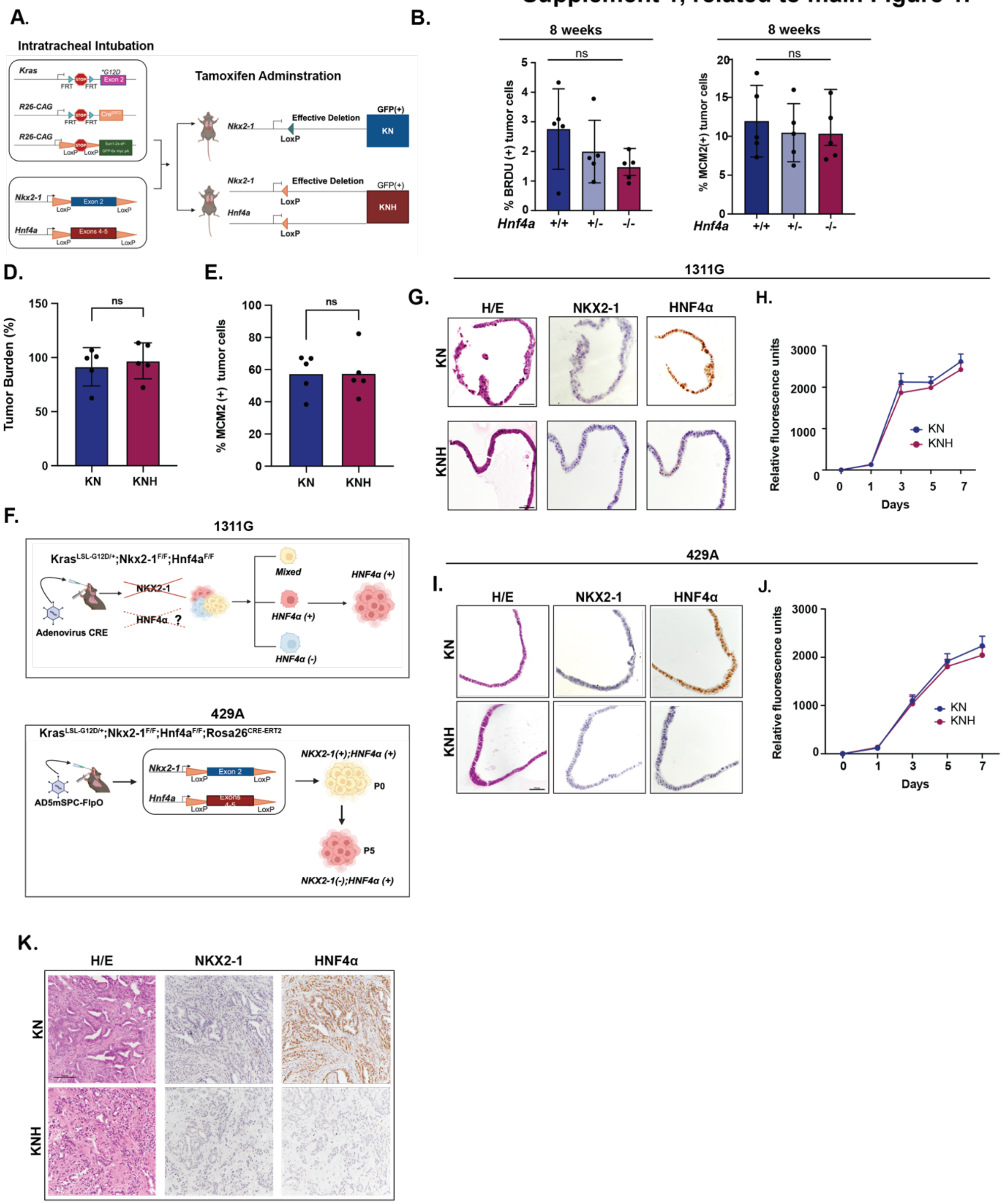

#### Supplement 1, related to Figure 1

**A**. Graphical representation for generating GEMM models of IMA.

**B-C**. Quantification of BrdU incorporation (**B**) and MCM2 expression (**C**) via IHC in KN and KNH GEMMs at 14 weeks post-tumor initiation. One-way ANOVA revealed no significant differences in BrdU incorporation (B, F (1, n) = 1.986, p = 0.1799, $R^2$ = 0.2486) or in MCM2 incorporation between genotypes (C, F (1, n) = 1.986, p = 0.7818, $R^2$ = 0.04019).

**D-E**. Mice were intubated with Ad5mSPC-FlpO virus and lungs were harvested one week after IP tamoxifen in KN (N=5) and KNH (N=5) GEMM for the quantification of overall tumor burden (**D**, unpaired t test; p = 0.4801)  and MCM2 expression via IHC staining of lung tissues (**E**, unpaired t test; p = 0.9837).

**F**. Schematic representation of organoid generation. Top: 1311G organoid line from a *Kras*$^{LSL\text{-}G12D/+}$*; Nkx2-1*$^{F/F}$*; Hnf4a*$^{F/F}$ mouse. Bottom: 429A organoid line from a *Kras*$^{FSF\text{-}G12D/+;}$ *Rosa26*$^{FSF\text{-}CreERT2}$*; Nkx2-1*$^{F/}$*F; Hnf4a*$^{F/F}$ mouse.

**G**. Representative images of IHC for NKX2-1 and HNF4α and H&E staining for 1311G KN and KNH organoids. Scale bar: 100 µm.

**H**. PrestoBlue viability assay measuring growth rates of KN and KNH 1311G organoids. Data shown represent one representative replicate from three biological replicates. Error bars indicate standard deviation (SD) of technical replicates.

**I**. Representative images of H&E staining and IHC for NKX2-1 and HNF4α of 1311G KN and KNH organoids. Scale bar: 100 µm.

**J**. PrestoBlue viability assay measuring growth rates of KN and KNH 429A organoids. Data shown represent one representative replicate from three biological replicates. Error bars indicate SD of technical replicates.

**K**. Representative images of H&E staining and IHC for NKX2-1 and HNF4α of subcutaneous tumors derived from allograft transplantation of the 429A IMA organoid line (KN and KNH) into NRG mice harvested at the endpoint. Scale bar: 250 μm.

#### Supplement 2, related to Main Figure 2

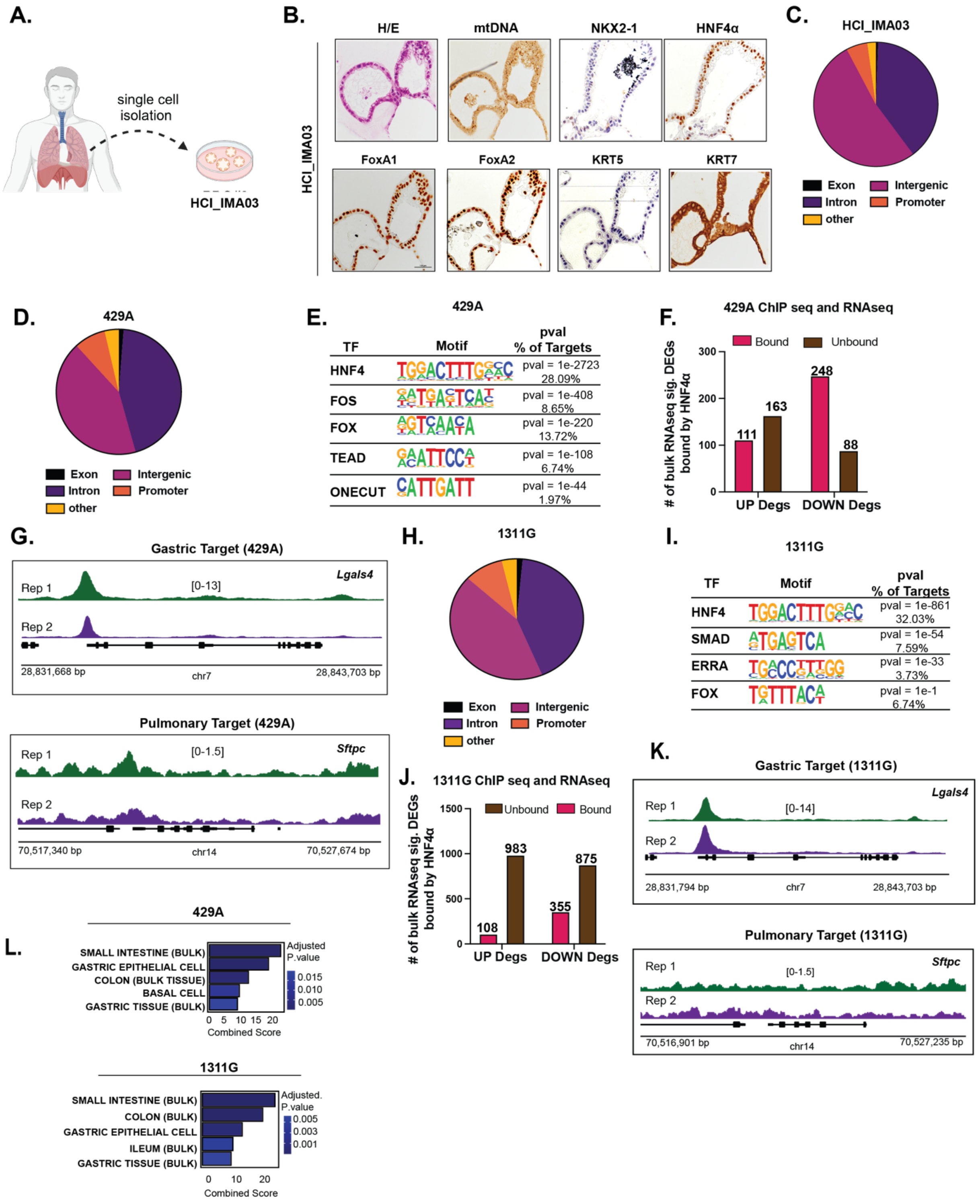

**Supplement 2, related to Figure 2.**

**A**. Schematic diagram of the derivation of HCI_IMA03 from an IMA tumor at the Huntsman Cancer Institute.

**B**. Representative images of H&E staining and IHC for NKX2-1, HNF4α, FoxA1, FoxA2, Cytokeratin 5 (KRT5), Cytokeratin 7 (KRT7), GKN1, human mitochondrion (mtDNA), and LGALS4 of HCI_IMA03. Scale bar: 100μm.

**C**. Genome-wide distribution of HNF4α ChIP-seq peaks across annotated genomic regions in HCI_IMA03.

**D**. Genome-wide distribution of HNF4α ChIP-seq peaks across annotated genomic regions in 429A.

**E**. HOMER motif enrichment analysis of HNF4α-bound peaks in 429A, ranked by enrichment score and p-value.

**F**. Bar plot showing overlap between HNF4α-bound regions and DEGs from bulk RNA-seq in 429A.

**G**. ChIP-seq tracks showing representative HNF4α binding at the gastric marker *Lgals4* and pulmonary marker *Sftpc* in 429A.

**H**. Genome-wide distribution of HNF4α ChIP-seq peaks across annotated genomic regions in 1311G.

**I**. HOMER motif enrichment analysis of HNF4α-bound peaks in 1311G.

**J**. Bar plot showing overlap between HNF4α-bound regions and DEGs from bulk RNA-seq in 1311G.

**K**. ChIP-seq tracks showing representative HNF4α binding at *Lgals4* and *Sftpc* in 1311G.

**L**. ENRICHR ARCHS4 tissue enrichment of genes annotated from HNF4α peaks in 1311G and 429A.

#### Supplement Fig 3, related to Main Figure 2.

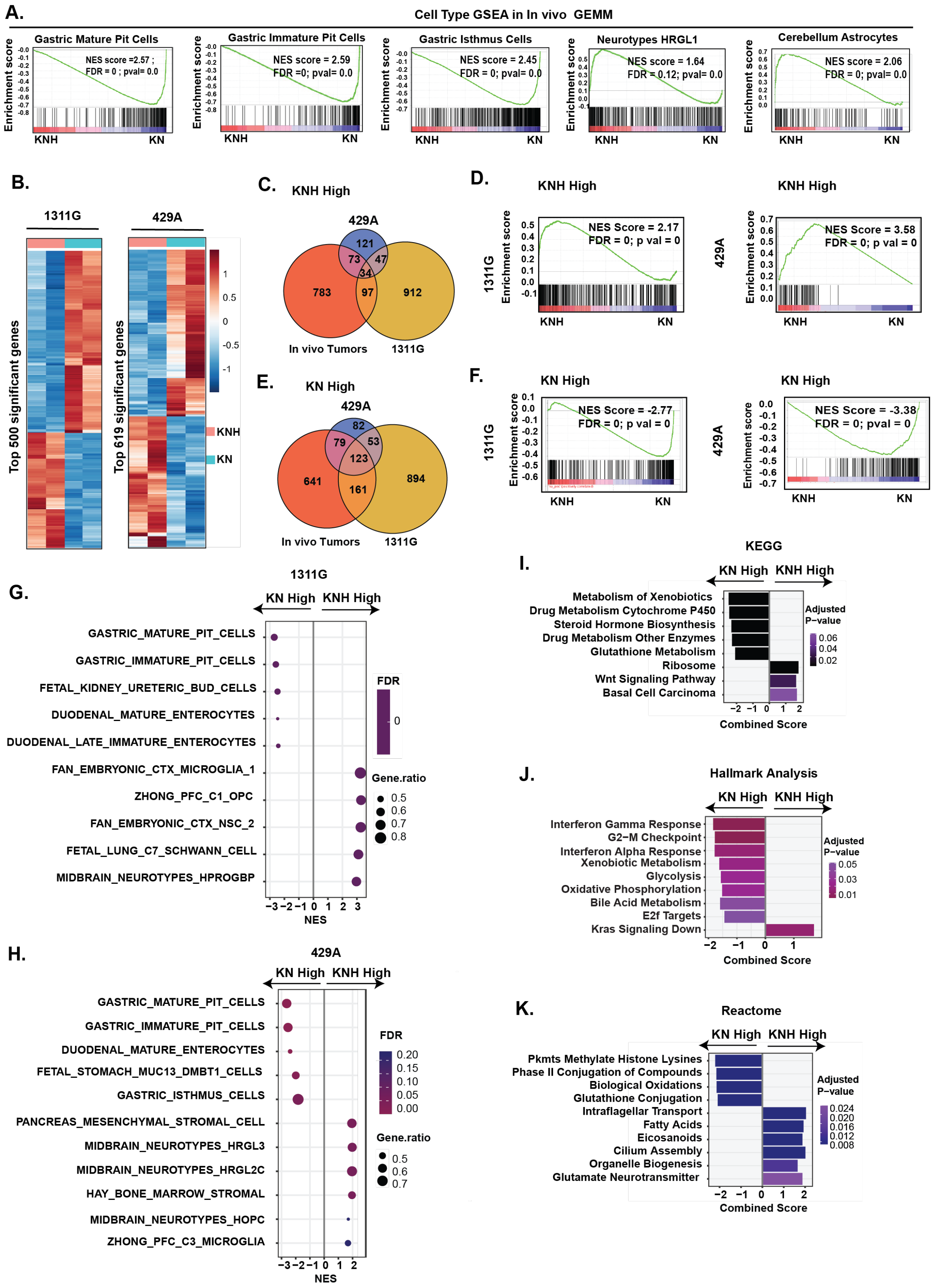

#### Supplement 3, related to Figure 2

**A**. GSEA plot of DEGs from bulk RNA-seq of KNH versus KN tumors, displaying gene signatures corresponding to gained and lost cell types (related to Fig. 2L). NES and FDR are indicated.

**B**. Heatmap of significant DEGs in isogenic IMA organoid lines: 1311G (Left: Top 500 genes) and 429A (right: Top 619 genes).

**C**. Venn diagram illustrating the overlap of KNH high DEGs identified in vivo and in the 1311G and 429A organoid lines.

**D**. GSEA plots of in vivo KNH high DEGs mapped onto the 1311G and 429A organoids. NES and FDR are indicated.

**E**. Venn diagram showing the overlap of KN High DEGs identified in vivo and in the organoid lines.

**F**. GSEA plots of in vivo KN High DEGs mapped onto the 1311G and 429A organoids.

**G-H**. C8 cell type enrichment analysis of DEGs in 1311G (**G**) and 429A (**H**) organoids.

**I**. KEGG pathway enrichment of shared DEGs from KNH versus KN tumors in vivo.

**J**. Hallmark pathway enrichment of DEGs from KNH versus KN tumors in vivo.

**K**. Reactome pathway enrichment of shared DEGs from KNH versus KN tumors in vivo.

### Supplement 4, related to Main Figure 3

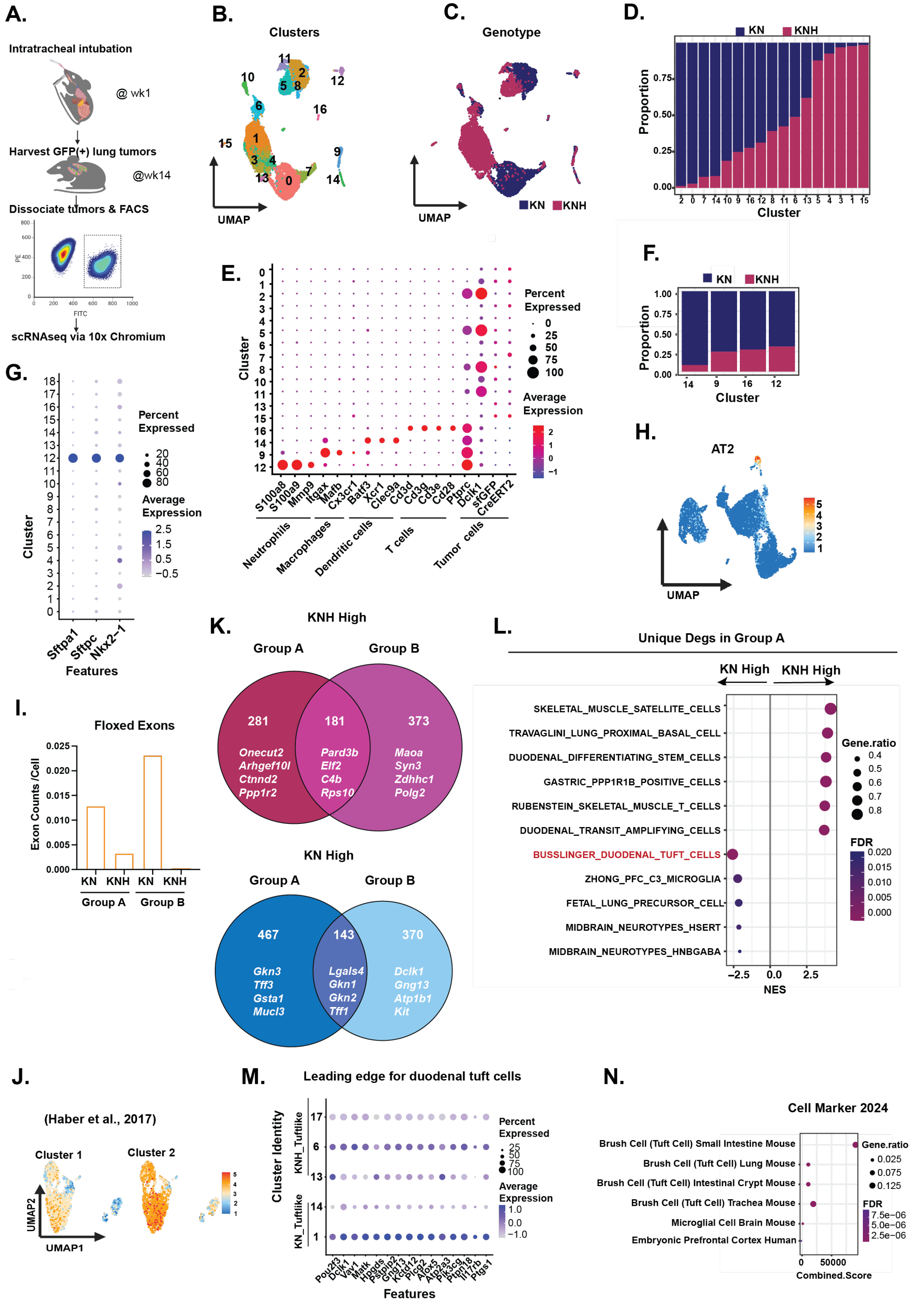

**Supplement 4, related to Figure 3**

**A**. Schematic of the experimental workflow for scRNA-seq analysis of GFP-positive tumors sorted from KN and KNH GEMMs.

**B**. UMAP of all captured cells from KN and KNH tumors (n = 2 mice per genotype, multiple tumors per mouse), colored by Seurat-defined clusters.

**C**. UMAP of the same dataset colored by genotype.

**D**. Proportion of KN and KNH cells across all captured cells.

**E**. Dot plot showing the expression of immune-related and tumor-associated marker genes across Seurat-defined clusters.

**F**. Proportion of KN and KNH cells within immune-enriched clusters that were excluded from downstream analysis.

**G**. Dot plot showing the expression of alveolar type 2 (AT2) markers, including *Sftpa1*, *Sftpc*, and *Nkx2-1*, across all bona fide tumor cells.

**H**. UMAPs showing gene module scores for AT2-like cells based on a previously defined gene set (PMID: 32707077).

**I**. Number of reads per cell aligning to *Hnf4a* floxed exons 4 and 5 in KN versus KNH tumor cells.

**J**. UMAP showing enrichment of Cluster 1 (neuronal-like) and Cluster 2 (immune-like) tuft-like cells in KN and KNH tumors within Group A (PMID: 29144463).

**K**. Venn diagrams showing overlapping DEGs between Group A and Group B. Top: KNH High DEGs. Bottom: KN High DEGs.

**L**. GSEA-based C8 cell type enrichment analysis of unique DEGs in tuft-like IMA cells comparing KN and KNH tumors.

**M**. Dot plot showing expression of leading-edge genes contributing to the downregulation of the duodenal tuft cell gene signature in KNH verses KN tumors.

**N**. ENRICHR cell type enrichment analysis of the leading-edge genes identified in panel M.

#### Supplement 5 related to Main Figure 3

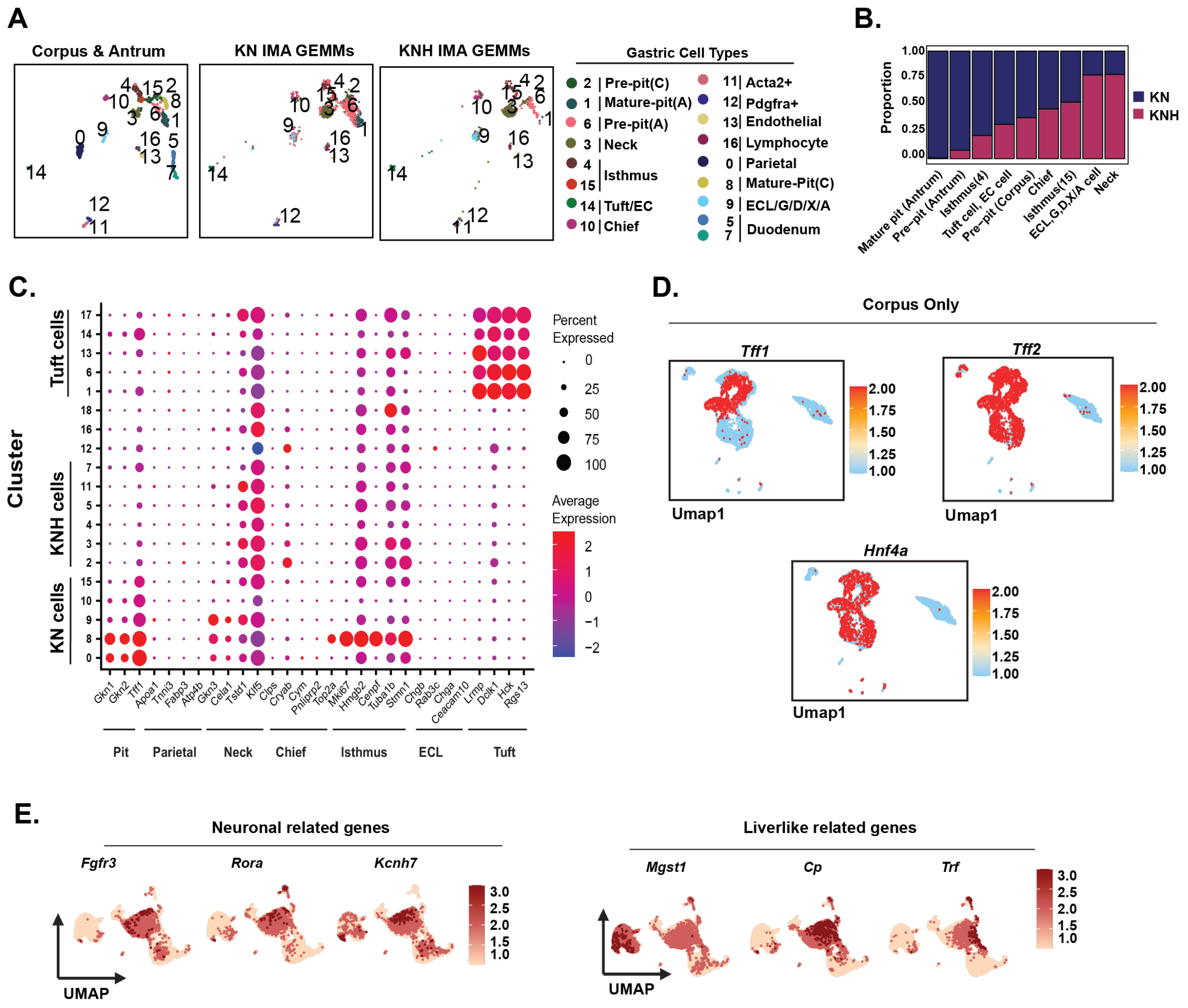

**<u>Supplement 5, related to Main Figure 3</u>**

**A**. UMAP of corpus and antrum dataset (PMID: 37386010) showing annotated gastric epithelial and stromal cell types (left), alongside UMAP of KN and KNH tumor cells mapped into the same reference space using Seurat label transfer (right).

**B**. Proportion of KN and KNH tumor cells mapping to each cluster in corpus and antrum dataset above.

**C**. Dot plot of representative marker genes across major gastric cell types in IMA.

**D**. Feature plots of *Tff1*, *Tff2* and *Hnf4a* mapped onto corpus only dataset (PMID: 37386010).

**E**. Feature plots showing expression of neuronal-associated genes (*Fgfr3, Rora and Kcnh7*) and liver-like genes (*Mgst1, Cp and Trf*) in IMA tumor cells.

**Supplement S6, related to Main Figure 4**

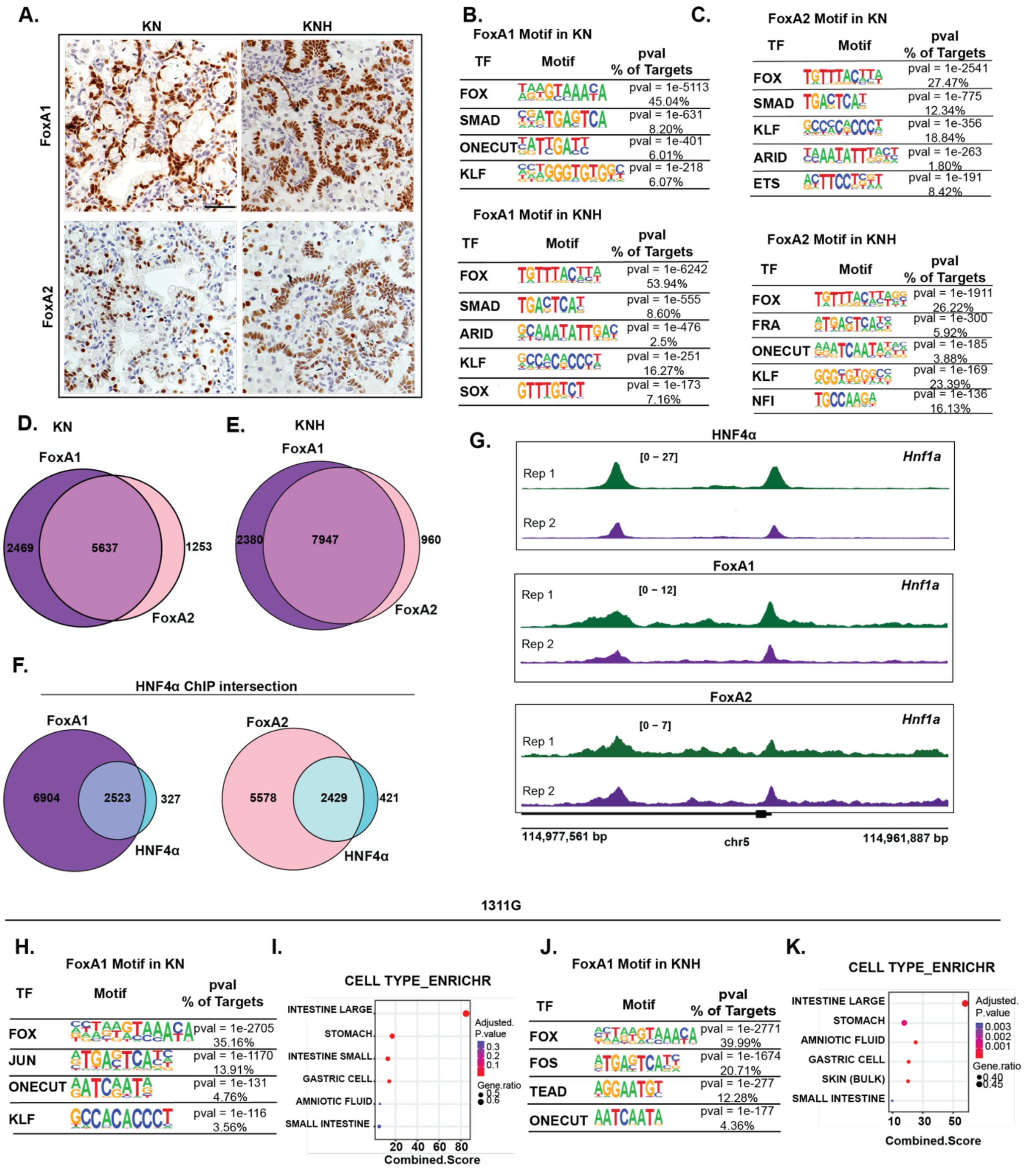

**Supplement 6, related to Main Figure 4**

**A.** Representative IHC staining for FoxA1 and FoxA2 in KN and KNH tumors at 14 weeks post-tumor initiation. Scale bar: 100 μm.

**B**. HOMER motif enrichment of FoxA1-bound peaks in KN and KNH tumors. Top: motifs enriched in KN-specific peaks. Bottom: motifs enriched in KNH-specific peaks.

**C**. HOMER motif enrichment of FoxA2-bound peaks in KN and KNH tumors. Top: motifs enriched in KN-specific peaks. Bottom: motifs enriched in KNH-specific peaks.

**D**. Overlap of annotated genes from FoxA1 and FoxA2 peaks in KN GEMMs.

**E**. Overlap of annotated genes from FoxA1 and FoxA2 peaks in KNH GEMMs.

**F**. Overlap of genes annotated from FoxA1 (left) or FoxA2 (right) peaks with those associated with HNF4α binding sites in KN tumors in vivo.

**G**. ChIP-seq tracks for FoxA1, FoxA2, and HNF4α in KN tumors showing co-occupancy at the gastric gene *Lgals4* (n = 2).

**H**. HOMER motif enrichment of FoxA1-bound peaks in 1311G KN organoids.

**I**. ENRICHR ARCHS4 tissue enrichment of genes annotated from FoxA1-bound peaks in 1311G KN.

**J**. HOMER motif enrichment of FoxA1-bound peaks in 1311G KNH organoids.

**K**. ENRICHR ARCHS4 tissue enrichment of genes annotated from FoxA1-bound peaks in 1311G KNH.

#### Supplement 7, related to Main Figure 4

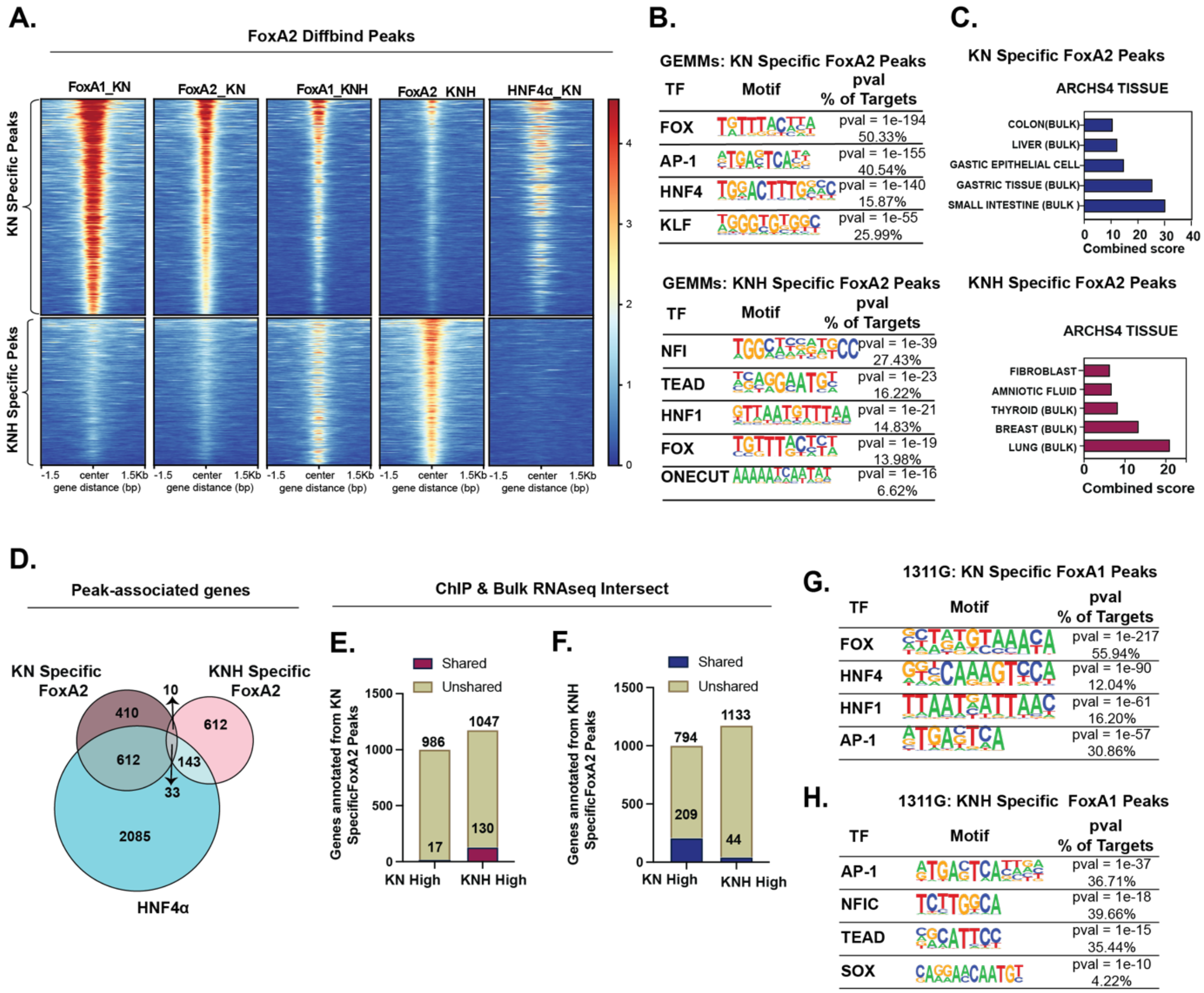

**Supplement 7, related to Figure 4**

**A.** Heatmap showing differential FoxA2 signal enrichment between KN and KNH tumors in vivo, identified using DiffBind (adjusted $p < 0.05$). Differential signal intensities were quantified over merged peak regions, defined as significant peaks (MACS2, adjusted $p < 0.05$) detected in at least one condition, spanning HNF4α, FoxA1, and FoxA2 in KN, and FoxA1 and FoxA2 in KNH.

**B.** HOMER motif enrichment of differential FoxA2 peaks. Left: motifs enriched in KN-specific peaks. Right: motifs enriched in KNH-specific peaks.

**C.** ENRICHR ARCHS4 tissue enrichment of genes annotated from differential FoxA2 peaks. Left: KN-specific. Right: KNH-specific.

**D.** Venn diagram showing the overlap between genes associated with differential FoxA2 peaks and those linked to HNF4α-bound regions in KN tumors in vivo.

**E.** Bar plot showing overlap between genes annotated from KN-specific FoxA2 peaks and DEGs from bulk RNA-seq.

**F.** Bar plot showing overlap between genes annotated from KNH-specific FoxA2 peaks and DEGs from bulk RNA-seq in vivo.

**G.** HOMER motif enrichment of differential FoxA1 peaks in 1311G KN organoids.

**H.** HOMER motif enrichment of differential FoxA1 peaks in 1311G KNH organoids.

#### Supplementary S8 for Main Figure 5

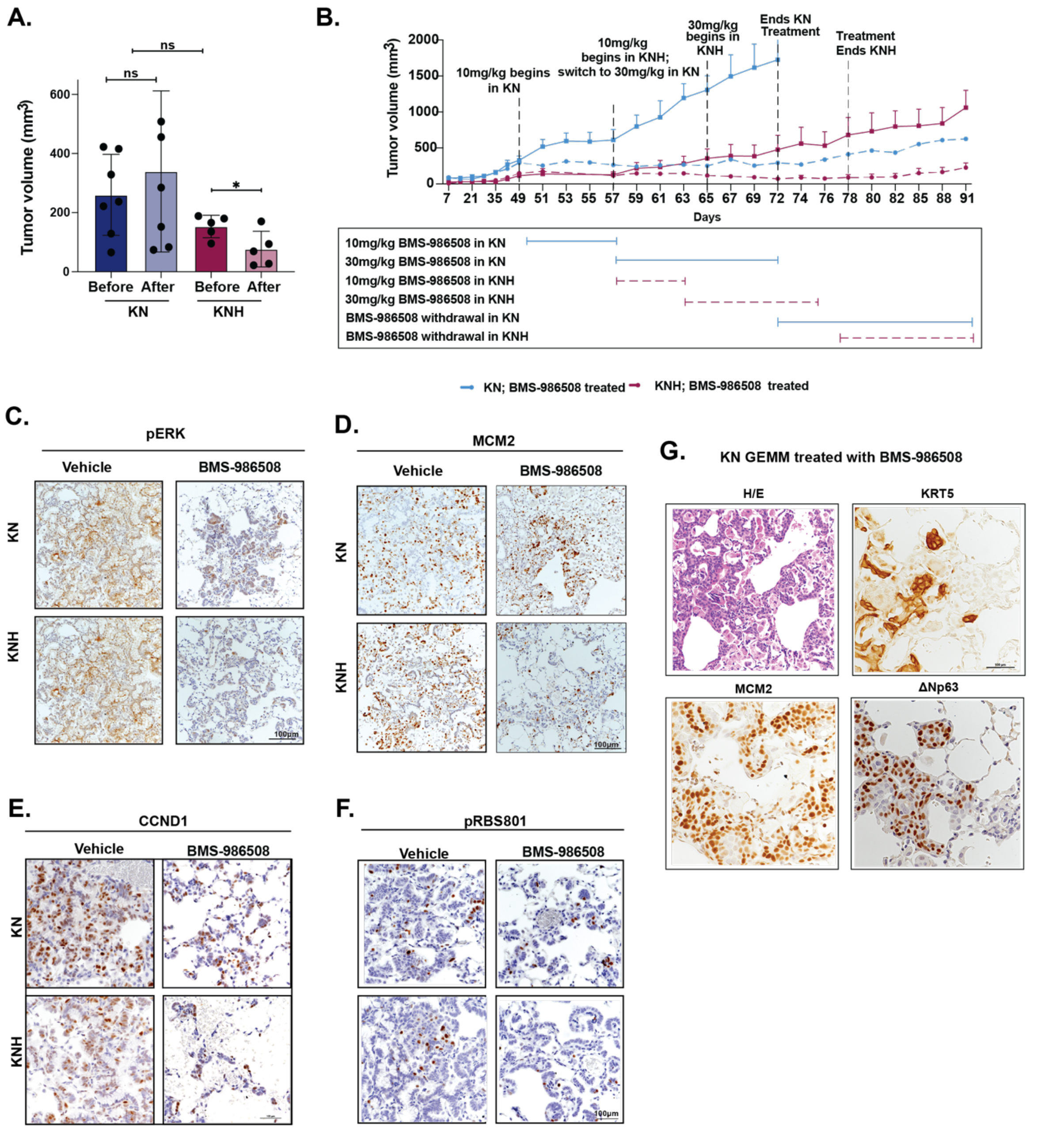

#### Supplement 8, related to Figure 5

**A.** Tumor volumes in individual mice bearing 1311G KN or KNH allografts before and after BMS-986508 treatment. Unpaired t-test: *p = 0.01239. Error bars represent SD from technical replicates.

**B.** Longitudinal tumor volume measurements from NSG mice implanted subcutaneously with 1311G KN or KNH organoids ($4 \times 10^5$ cells). Treatment began at ~150 $mm^3$. Mice received 10 mg/kg BMS-986508 or vehicle for 7 days, then 30 mg/kg for 14 days, followed by a weaning period. Error bars represent SEM from technical replicates.

**C.** Representative IHC images for pERK in tumors from GEMMs (KN and KNH) treated with 30 mg/kg BMS-986508 or vehicle for 14 days (weeks 12-14 post-initiation). Scale bar: 100 µm.

**D-F**. Representative IHC images of proliferation and cell cycle markers in KN and KNH GEMMs treated in vivo with 30 mg/kg BMS-986508 or vehicle for 14 days. Shown are MCM2 (**D**), CCND1 (**E**), and pRBS807 (**F**). Scale bar: 100 µm.

**G.** Representative IHC images for MCM2, CK5, ΔNp63, and H&E staining in tumors from a KN mouse treated in vivo with 30 mg/kg BMS-986508 or vehicle for 14 days. Scale bar: 100 µm.

#### Supplement S9, related to Main Figure 6

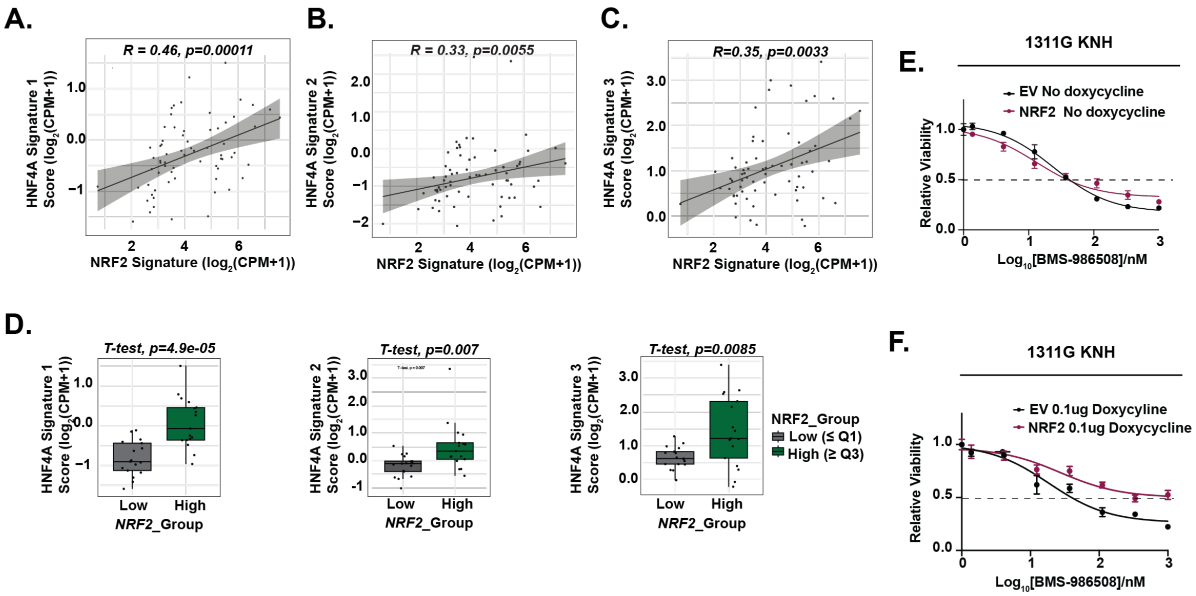

**Supplement 9, related to for Figure 6**

**A-C.** Spearman correlations between NRF2 activity and HNF4A signature scores across 68 KRAS-mutant NSCLC tumors from the KRYSTAL-1 dataset. NRF2 activity was quantified using a published NRF2 signature. HNF4A activity was assessed using three independent gene signatures: Signature 1 (**A**), Signature 2 (**B**), and Signature 3 (**C**). Each dot represents a tumor. Gray regression lines and 95% confidence intervals are shown. Spearman correlation coefficients (*R*) and *p*-values are indicated.

**D.** Boxplots comparing HNF4A signature 1, 2 and 3 activity scores between *NRF2*_Low and *NRF2*_High tumors. All three signatures were significantly elevated in the NRF2_High group. p-values were determined using unpaired two-tailed Student's *t*-test.

**E-F.** NRF2 re-expression increases BMS-986508 resistance in 1311G KNH organoids. Organoids expressing empty vector (EV - **E**) or DOX-inducible NRF2 (**F**) were treated with BMS-986508 for 72 h in the absence (ND) or presence of 0.1 µg/mL DOX. $IC_{50}$ increased from 2.5 nM (EV) to 14.2 nM (NRF2 + DOX), indicating partial rescue of sensitivity. Data represent one of three independent experiments. Error bars represent SEM from technical replicates.

**Supplement S10, related to Main Figure 6**

**A.**

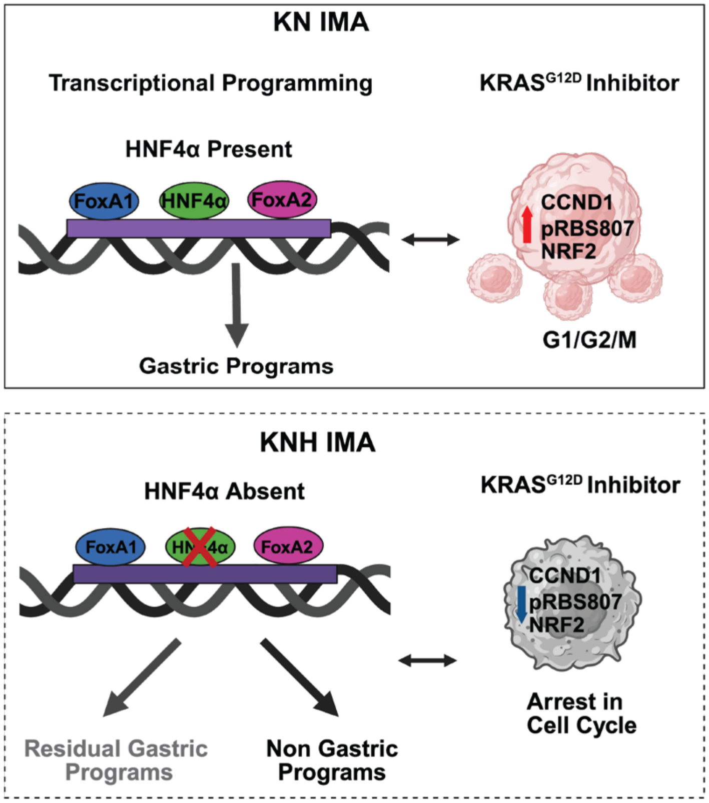

**Supplement 10, related to for Figure 6**

A. Overall working model depicting the role of HNF4α as a key regulator of gastric identity programs and primary response to $KRAS^{G12D}$ inhibition in IMA.
